# Failed remyelination of the non-human primate optic nerve leads to axon degeneration, retinal damages and visual dysfunction

**DOI:** 10.1101/2022.01.11.475669

**Authors:** Nadège Sarrazin, Estelle Chavret-Reculon, Corinne Bachelin, Mehdi Felfli, Rafik Arab, Sophie Gilardeau, Elena Brazhnikova, Elisabeth Dubus, Lydia Yaha-Cherif, Jean Lorenceau, Serge Picaud, Serge Rosolen, Pierre Moissonnier, Pierre Pouget, Anne Baron-Van Evercooren

## Abstract

White matter disorders of the CNS such as MS, lead to failure of nerve conduction and long-lasting neurological disabilities affecting a variety of sensory and motor systems including vision. While most disease-modifying therapies target the immune and inflammatory response, the promotion of remyelination has become a new therapeutic avenue, to prevent neuronal degeneration and promote recovery. Most of these strategies are developed in short-lived rodent models of demyelination, which spontaneously repair and do not reflect the size, organization, and biology of the human CNS. Thus, well-defined non-human primate models are required to efficiently advance therapeutic approaches for patients. Here, we followed the consequence of long-term toxin-induced demyelination of the macaque optic nerve on remyelination and axon preservation, as well as its impact on visual functions. Findings from oculo-motor behavior, ophthalmic examination, electrophysiology, and retinal imaging indicate visual impairment involving the optic nerve and retina. These visual dysfunctions fully correlated at the anatomical level, with sustained optic nerve demyelination, axonal degeneration, and alterations of the inner retinal layers. This non-human primate model of chronic optic nerve demyelination associated with axonal degeneration and visual dysfunction, recapitulates several key features of MS lesions and should be instrumental in providing the missing link to translate emerging repair pro-myelinating/neuroprotective therapies to the clinic for myelin disorders such as MS.

**Significance Statement:** Promotion of remyelination has become a new therapeutic avenue, to prevent neuronal degeneration and promote recovery in white matter diseases such as MS. To date most of these strategies are developed in short-lived rodent models of demyelination, which spontaneously repair. Well-defined non-human primate models closer to man would allow to efficiently advance therapeutic approaches. Here we present a non-human primate model of optic nerve demyelination that recapitulates several features of MS lesions. The model leads to failed remyelination, associated with progressive axonal degeneration and visual dysfunction, thus providing the missing link to translate emerging pre-clinical therapies to the clinic for myelin disorders such as MS.

## Introduction

White matter disorders are a large group of neurological diseases of various origins. Those affecting the central nervous system (CNS) such as multiple sclerosis (MS) lead to failure of nerve conduction, axon degeneration, and result in long-lasting neurological disabilities and tissue atrophy (1). The loss of myelin and healthy axons are believed to be responsible for irreversible damages, which affect a variety of sensory and motor systems including vision. In MS, seventy percent of patients are affected with optic neuritis. It can manifest in an acute episode with decreased vision that can recover over several weeks in the majority of patients while permanent visual symptoms persist in 40-60% of patients (2, 3). Chronic optic neuritis can lead to significant optic nerve atrophy and retinal alterations affecting mainly the retinal inner layers including the retinal nerve fiber and ganglion cell layers (4). Several visual assays including visual fields (VF) (5), pupillary responses to luminance and color (PLR) (6), electroretinograms (ERG) (7), optical coherence tomography (OCT) (4, 8), and visual evoked potential (VEP) (9–11) are routinely performed to assess non-invasively the anatomical and/or electrophysiological perturbations of visual functions in MS. While functional recovery was reported in some patients (9), the lack of anatomical-electrophysiological correlation has prevented to attribute directly these improvements to remyelination or other regenerative processes.

Animal models of demyelination induced by toxins such as lyso-phosphatidylcholine (LPC) are suitable to study the mechanisms of demyelination-remyelination and develop approaches aiming at promoting CNS re-myelination as they show little inflammation and therefore, provide means to assay directly the effect of a therapy on remyelination. However, most of these models are developed in short-lived rodents and spontaneously repair, thus lacking the long-lasting progressive degenerative disease context of MS. Besides, these models do not reflect the size, or complex organization of the human primate CNS (12). They do not inform on the biology of primate cells, which differs from rodents (13, 14), nor on the security, toxicity, and long-term efficacy of cell- or compound-based pro-myelinating/neuroprotective therapies. Thus, experiments in long-lived non-human primates appear an essential step towards clinical trials.

While promoting remyelination may prevent axon degeneration, only few pro-myelination strategies have been translated to the clinic (15,17). One of the roadblocks is the absence of studies addressing the clinical benefit of pro-myelination approaches that could be applied to the clinic (17). A positive correlation between changes in VEP parameters and the degree of demyelination/remyelination was established in rodents (18–21), cats (22), and dogs (23), and exploited successfully to follow pro-myelination therapies in rodents (24, 25). OCT has been used to identify loss of optic nerve and retinal damages in animal models of myelin disorders as well (23, 26). While used seldomly in non-human primates (27), none of these clinical assays were exploited to monitor the impact of optic nerve demyelination in non-human primates.

We previously demonstrated that LPC injection in the macaque optic nerve induced demyelination with fair axon preservation but little remyelination up to 2 months post-demyelination (28). Taking advantage of the fact that non-human primates are long-lived and are able to perform several tasks awake such as humans, we questioned whether this model could be used to follow the consequence of long-term demyelination on axon preservation, and whether multimodal non-invasive assays such as VF, VEP, OCT and PLR could be instrumental to follow/predict the functional and anatomical outcome of optic nerve demyelination. Using multidisciplinary approaches, we provide compelling evidences that LPC-induced demyelination of the macaque optic nerve leads to modified VF, VEP, PLR, and altered inner retinal layers but preserved photoreceptors based on OCT and ERG. These clinical and functional anomalies were correlated at the histological level with failed remyelination and progressive optic nerve axon loss, followed by neuronal and fiber loss of the inner retinal layers. The post-mortem validation of OCT, VEP and PLR as pertinent markers of optic nerve demyelination/degeneration could further help the translation of therapeutic strategies towards the clinic for myelin diseases associated with long-term demyelination of the optic nerve.

## Results

The impact of LPC-induced demyelination of the optic nerve on visual functions was studied using psycho-physic (VF), electro-physiologic (VEP, ERG), physiologic (PLR) and anatomic (OCT) tools. As saline and LPC injections could not be performed in the same animal for ethical reasons, we compared responses of the left eye corresponding to the LPC injected optic nerve (LPC-ON) with that of the right fellow eye corresponding to the non-injected optic nerve (C-ON). Data were further correlated with post-mortem analysis of the optic nerves and associated retinas of the same subjects (Table S1).

### Altered visual field sensitivity

Measuring VF sensitivity is long and challenging with awake monkeys. Therefore, we developed an in-house method relying on analyzing the latencies of saccades toward targets presented at series of central and eccentric locations. Indeed, visual defects resulting in unseen or poorly seen targets are expected to entail long saccadic latencies, no saccade at all, or an erratic oculomotor behavior. A Gaze-contingent display that triggers the successive presentations of the visual targets was used to analyze the saccadic latencies (the time for each eye to reach the new target in monocular and binocular conditions). Although this method does not directly evaluate visual field sensitivity, it proved useful to identify functional visual field defects consecutive to the LPC induced demyelination (Fig. 1A, Fig. S1A, Table S2). Statistical analysis showed a significant effect of time (*P*=1.23e-10***) as well as a significant interaction time:eye (*P*<2e-16***). While global mean saccade latencies between C-ON and LPC-ON did not differ significantly pre-lesion, they changed only slightly post-lesion for the C-ON compared to the LPC-ON corresponding eye. Post-hoc analysis per time point showed a significant increase in the mean saccade latencies for the LPC-ON compared to the C-ON for each time point post-LPC. As the latencies measured herein corresponded to the time between the current fixation and the fixation of a next target, these LPC-ON long latencies presumably reflect both the time required to search for the target with residual visual sensitivity as well as the increased latency due to optic nerve degeneration. The large latency differences indicate that the LPC-induced lesion triggered important visual deficits, suggestive of retinal modifications.

**Fig 1.**
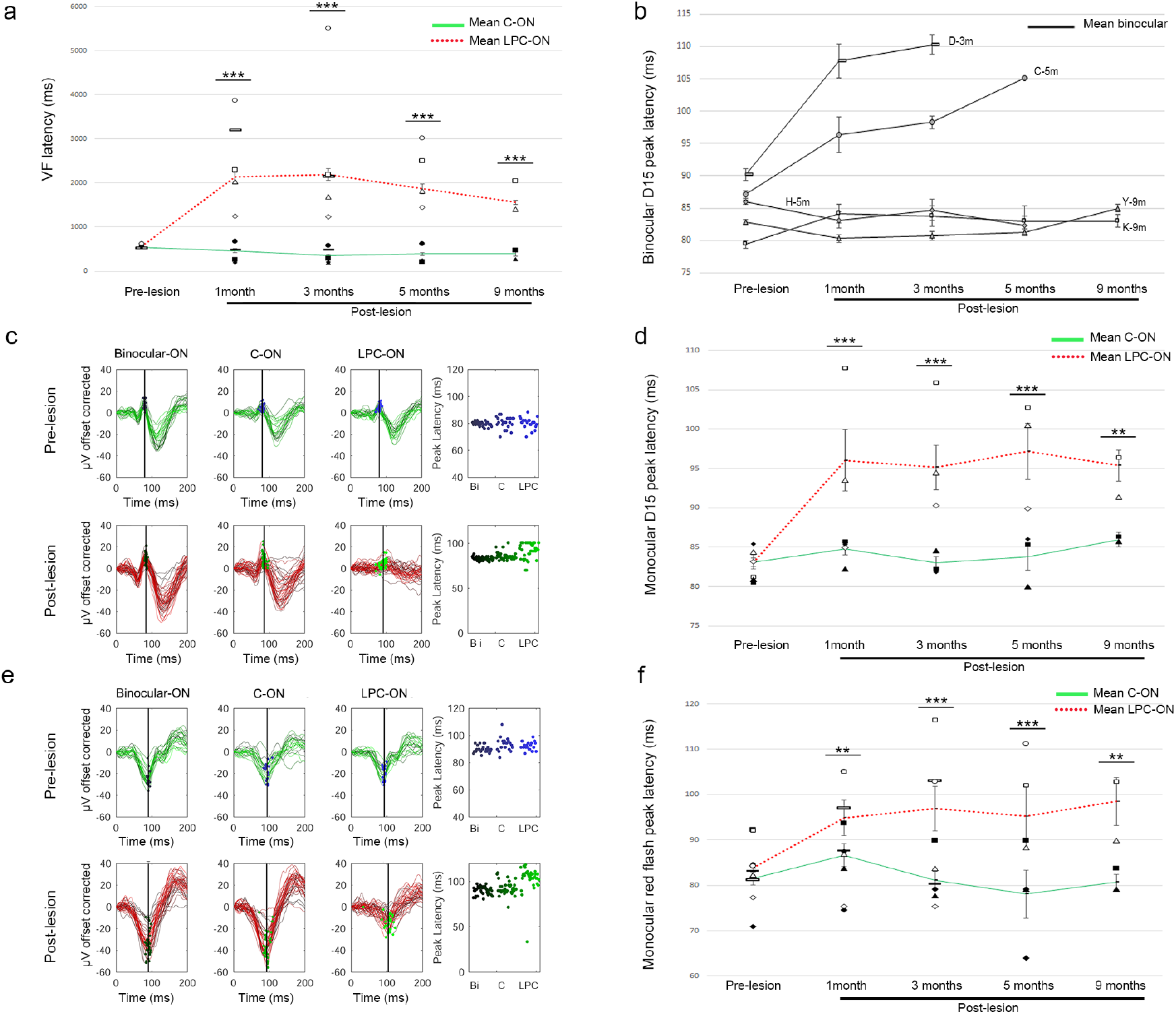
Longitudinal evolution of VF and VEPs in response to optic nerve LPC-induced demyelination. **(A)** Diagram of VF latencies of C-ON and LPC-ON corresponding eyes. Differences between pre- and post-LPC are not significant for C-ON (n=5, *P*=0.373) and LPC-ON (n=5, *P*<0.662). Differences between eyes are significant from 1 month and 3 months, n=5; 5 months n=4; 9 months, n=2, *P*<0.001). **(B)** Diagram of D15 checkerboard latencies in binocular conditions (1-, 3 months, n=5; 5 months n=4; 9 months, n=2). Two out of 5 subjects have altered D15 latencies in binocular conditions. **(C)** Representative curves of D15 checkerboard, P80 peak in binocular and monocular conditions for one subject. **(D)** Diagram of D15 checkerboard latencies in monocular conditions. Differences between pre- and post-LPC are significant for LPC-ON only (n=3, *P*=0.056). Differences between eyes are significant at 1, 3, 5 (n=3, *P*<0.0001) and 9 (n=2, *P*=0.0053) months. **(E)** Representative curves of red flashes, N2 peak in the binocular and monocular conditions for one subject. **(F)** Diagram of red flashes N2 latencies in monocular conditions for C-ON and LPC-ON. Differences between pre- and post-LPC are significant for LPC-ON only (n=3, *P*=0.049). Differences are significant at 1- (n=3, *P*=0.0025), 3- (n=3, *P*<0.0001), 5- (n=3, *P*<0.0003) and 9 (n=2, *P*=0.0092) months post-LPC. In **D** and **F** full symbols represent base-line data and empty symbols post-demyelination data; subject correspondence is as follows: ■ or □ K-9m; ● or ○ C5-m; ▐ or ▯ D-3m; ◆ or ◇ H-5m; ▲ or Δ Y-9m. Asterisks (*) indiquate the significant difference between C-ON and LPC-ON for a given time point.

### Delayed visual evoked potential latencies

To ascertain that LPC entailed visual deficits, and not the oculomotor system, we further used VEP as an objective measure of the visual pathway function. In particular, we used flash VEP and pattern reversal checkerboard-evoked potentials on awake animals to mimic as closely as possible the visual assays performed on human subjects. The control macaque VEP recorded from the scalp was characterized by wavelets that preceded a major peak around 80ms. While changes in waveforms and amplitudes, were observed for all subjects in response to D15, D30, and D60 checkerboard and were quite heterogeneous, we focused our analysis on P80 latency responses for D15 and D30. First, we examined P80 latency responses to the D15 checkerboard in binocular conditions at 1-, 3-, 5-, and 9 months post-LPC. While 3 animals showed stable binocular responses before and post-LPC, 2 of them had disturbed binocular responses suggesting that their C-ON was unable to compensate for the LPC-ON. (Fig. 1B). Next, we analyzed D15 VEP responses for those animals with stable binocular responses in monocular condition (Fig. 1C, 1D). Statistical analysis showed a significant effect of time (*P*=0.0104*) as well as a significant interaction time:eye (*P*=1.05e-05***). Post-hoc analysis (Table S3) showed that pre-lesion average VEP latencies did not differ significantly between both eyes. While VEP latencies remained stable post-lesion for C-ON, they increased significantly post-lesion for the LPC-ON corresponding eye. Moreover, analysis at each time point, indicated a statistically prolonged latency for the LPC-ON corresponding eye relative to C-ON and as early as 1month post-LPC. Latency differences were also observed for responses to the D30 checkerboard (Fig. S1B, Table S3).

Abnormal N2 latency responses to red (Fig. 1E, F, Table S3) and blue flashes (Fig. S1C, Table S3) were also analyzed. A significant effect of time (*P*=0.000128***) as well as a significant time:eye interaction (*P*=3.92e-07***) was found for red flashes. Post-hoc analysis indicated that pre-lesion average VEP latencies responses to red flashes did not differ significantly between CON and LPC-ON. However, they increased markedly post-LPC for the LPC-ON corresponding eye, compared to C-ON. Moreover, post-hoc analysis for each time-point showed that N2 latencies were prolonged for the LPC-ON corresponding eye over the C-ON corresponding fellow eye as off 1month post-LPC for all 5 animals with significant mean value differences for all time points post-LPC. Statistical differences were also observed with blue flashes at 1- and 3 months post-LPC, but not at later times. Thus, analysis of VEPs (checkerboard and flashes) indicates that axon and their cell bodies are involved in visual impairment.

### Changes in pupillary light reflex responses

PLR involves a short pathway, from the retina to the ciliary ganglions innervating the iris sphincter, with relays in the PON and the Edinger-Westphal nucleus. Given that PLR is altered in optic neuritis, and used as a measure of optic nerve dysfunction, PLR responses were measured before and after lesion. First, we analyzed PLR responses to increasing intensities of blue and decreasing intensities of red flashes prior LPC injection. We focused on the C2 negative peak as a measure of pupillary maximal constriction, followed by PIPR (blue flashes only) as a measure of post-illumination pupillary reflex or partial recovery to a normal pupil state. We constructed an irradiance-response curve to 1-sec light exposure for each animal in non-lesion condition (Fig. 2A). Pupil diameter changed dynamically over the entire range of irradiance, with optimal constriction (C2) from 2.01 logCd/m^2^ and PIPR responses detectable in blue conditions only. Next, we compared the pupillary responses of the C-ON and LPC-ON corresponding eyes over time at 2.01logCd/m^2^. C2 (Fig. 2D) and PIPR (Fig. 2C) values expressed in percentage of baseline in non-stimulated conditions, did not differ significantly pre-LPC between C-ON and LPC-ON in blue or red stimulation (Table S4). Moreover, lesion induction in the LPC-ON had no impact on the C-ON since C2 and PIPR values of the control eye did not change pre- and post-LPC in response to blue and red stimulations. Statistical analysis of pupillary constriction based on C2 values showed a significant effect of time, as well as a significant interaction time:eye for blue (time, *P*=0.001550; time:eye, *P*=0.002528) and red flashes (time, *P*=0.004032, time:eye, *P*=4.546e-05). Pupillary constriction decreased at all times post-LPC in the LPC-ON in response to blue flashes and red flashes. PIPR responses of LPC-ON corresponding eye, detectable in blue conditions only, decreased dramatically at all time points post-LPC. Thus, the lesion affected both the maximum constriction and post-illumination pupillary light reflex.

**Fig 2.**
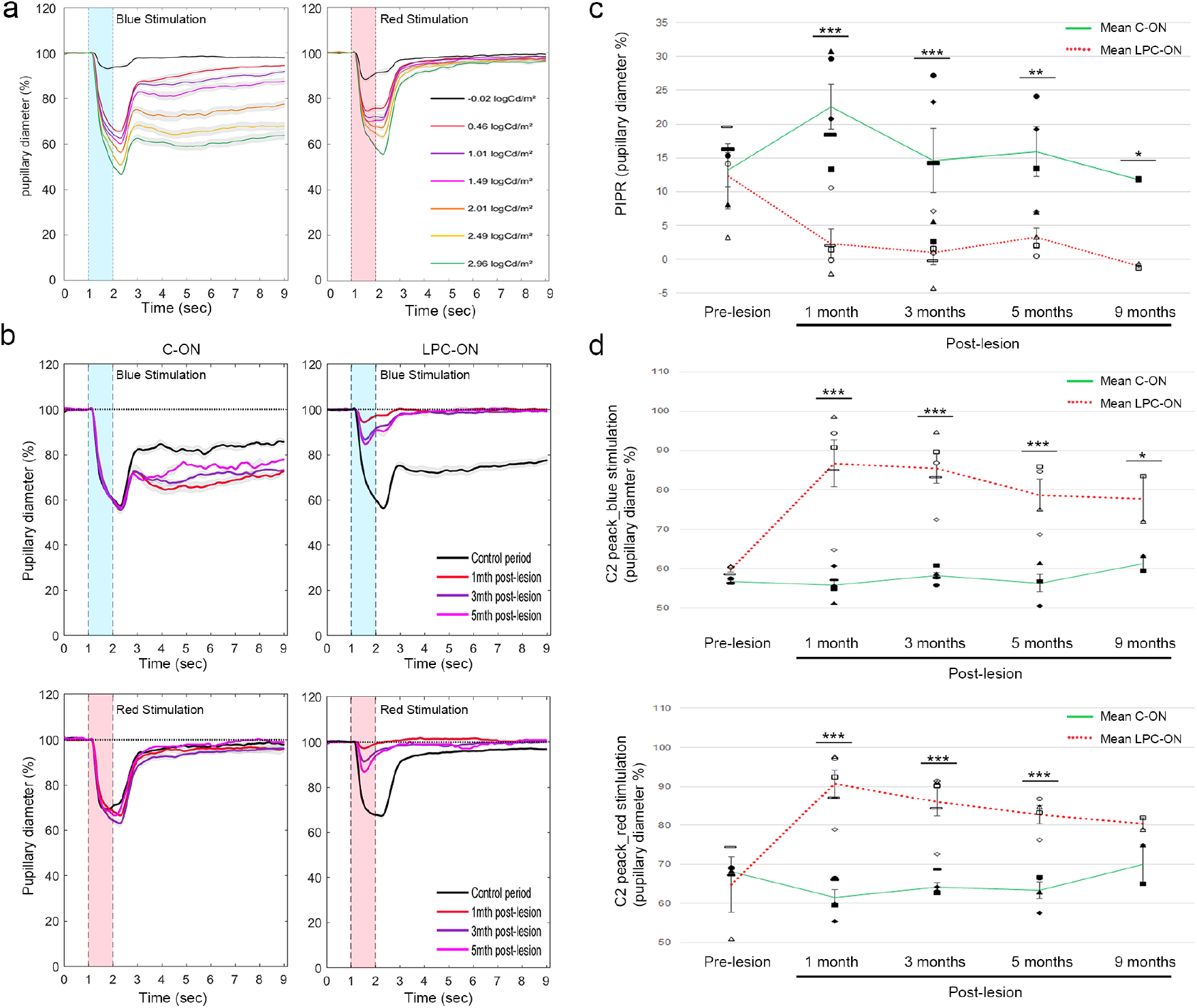
Longitudinal lesion-induced changes in PLR. **(A)** Pupillary responses after blue or red flashes at different intensities. **(B)** Representative pupillary responses to 2.01 log Cd/m^2^ post-LPC after blue (top) and red (bottom) stimulation. **(C)** Diagram of PIPR in response to blue stimulation. Differences between pre- and post-LPC are significant for LPC-ON only at 1-(n=5, *P<0,00001*), 3- (n=5, *P<0,0001*), 5- (n=3, *P<0,001*) and 9 (n=2, *P=0,03*) months, *P*). **(D)** Diagram of C2 peak in response to blue (top) and red (bottom) stimulation. Differences between pre- and post-LPC are significant (for LPC-ON only. Differences between eyes are significant at 1-, 3- (n=5, *P<0,00001*), 5 (n=3, *P<0,00001*) and 9 (n=2, *P<0,01*) months in blue condition. In red conditions, differences between eyes are significant at 1-, 3- (n=5, *P<0,001*), and 5- (n=3, *P<0,001*) months post LPC. In **B** and **C** full symbols represent base-line data and empty symbols post-demyelination data; subject correspondence is as follows: ■ or □ K-9m; ● or ○ C5-m; ▐ or ▯ D-3m; ◆ or ◇ H-5m; ▲ or Δ Y-9m. Asterisks (*) indiquate the significant difference between C-ON and LPC-ON for a given time point.

### Preservation of electroretinograms

To further investigate whether LPC-induced demyelination of the optic nerve did affect the most outer retinal layers, we performed ERG before and after LPC injection and compared the ERG responses of the LPC injected side over the non-injected side. Wave parameter’s mean values (± SD) before (n=3) and after LPC injection of the optic nerve (n=4) are presented in Table S5 and illustrated Fig. 3A. For all animals, each eye and each condition, ERG responses were discernable from the background noise. Before LPC, in all stimulation conditions, ANOVA analysis (*P*<0.05) did not yield any difference between the right CON and left LPC-ON corresponding eyes. Amplitudes slightly decreased post-LPC compared to values before LPC in both eyes. Moreover, mixed ERG waves and normal photopic and scotopic A and B waves were unchanged after LPC-induced demyelination compared to the internal control right eye and pre-LPC left eye. These observations predict that the photoreceptors (cone and rods) and their outputs are not affected by the LPC-induced demyelination.

**Fig 3.**
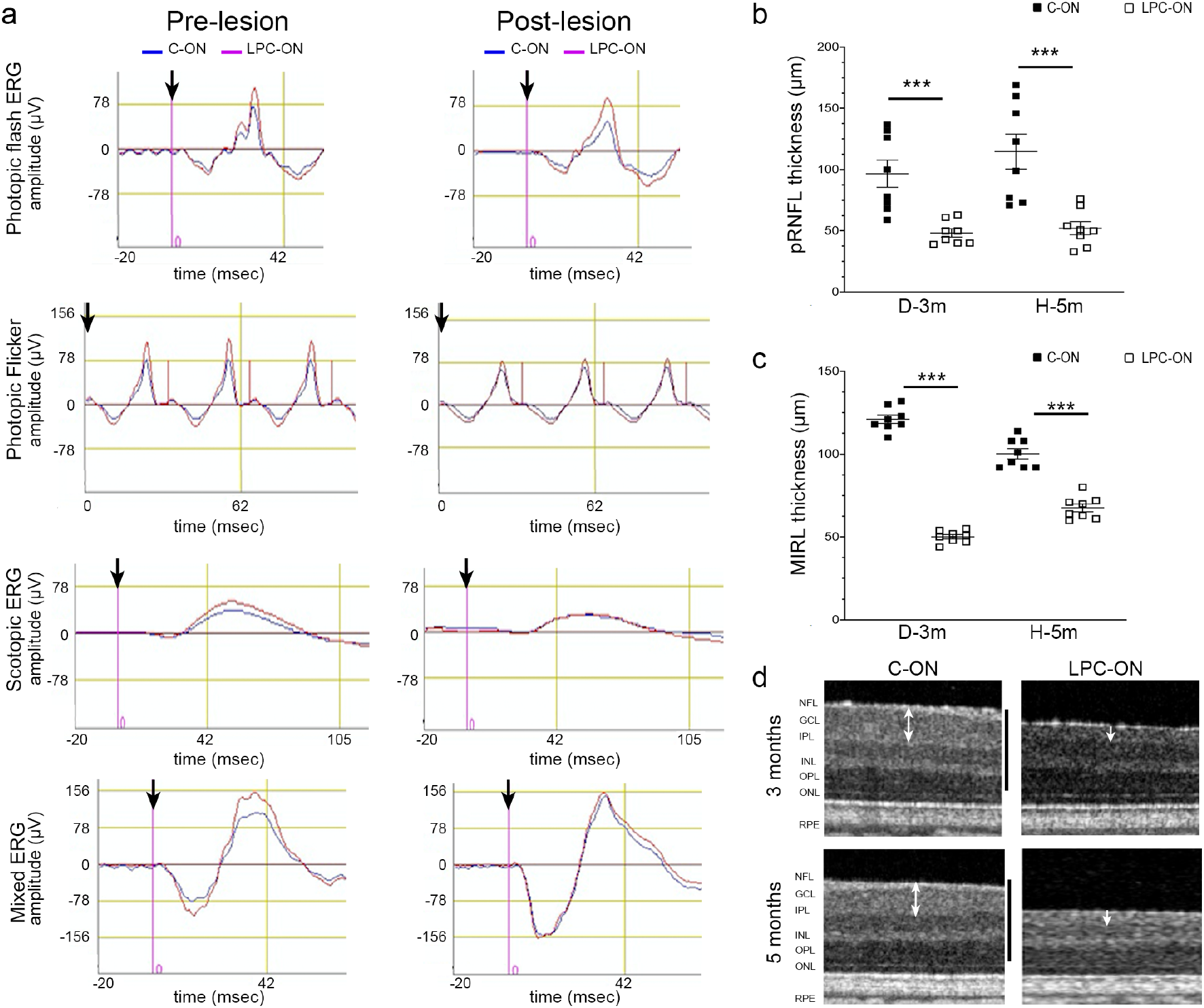
Evaluation of lesion-induced retinal changes by ERG and OCT. **(A)** Representative ERG waves showing Mixed ERG waves and normal photopic and scotopic A and B waves, which are unchanged post-LPC (Red) compared to the internal C-ON corresponding fellow eye (blue), and pre LPC lesion LPC-ON corresponding eye. The black arrow represents the flash onset. **(B, C)** The average thickness of **(B)** peri-papillary retinal nerve fiber layer (pRNFL) and **(C)** Macular inner layer (MIRL) thickness of retinas corresponding to C-ON (full columns) and LPC-ON (empty columns) at 3- (n=1) and 5- (n=1) months post-LPC. **(D)** MIRL of retinas corresponding to the CON, and LPC-ON at 3- and 5 months post-LPC injection. White arrows point to macular inner layer thickness. Data are the mean ± SEM for each monkey, t-Test \*\*\**P*<0.001. Scale bars: (C, D) 250μm.

### Optical coherence tomography imaging changes

Finally, we used OCT to assess the effects of LPC on the peri-papillary retinal nerve fiber layer (pRNFL) thickness and the macular inner retinal layer (MIRL) thickness. OCT measurements were performed before euthanasia at 3- (n=1) and 5 (n=1) months post-LPC injection of the ON.

The average pRNFL thickness measured by RNFL 3.45 scanning post-LPC was 50.19±1.94 μm for the LPC-ON corresponding eye, compared to the average pRNFL thickness of 105.69±9.06 μm for the internal C-ON corresponding eye (n=2) respectively, the difference being of 53% (D-3m: *P*=0, 0009; H-5m: *P*=0,001) (Fig. 3B). The macular inner layer (MIRL) was then automatically calculated on the superior, nasal, inferior, and temporal quadrants in the ETDRS - type areas by the Retina map scan pattern (Fig. 3C). The average MIRL thickness at 3- and 5 months after injection from two LPC-ON corresponding eyes was 58.53±9.1 μm compared to 110.69 ±10.4 μm in the two C-ON, the difference of MIRL thickness being of 52% (D-3m: *P*=5,4e-13; H-5m: *P*=9e-07). Analysis of MIRL thickness at 3- and 5 months post-LPC, showed a thickness decline of all measured macular areas (Fig. 3D). Altogether, the OCT data indicate that LPC induced demyelination of the optic nerve worsens in time and space, the retinal thickness of the LPC-ON corresponding eye without affecting the retina of the C-ON corresponding fellow eye.

The assessment of the optic nerve head (ONH) parameters with the Glaucoma scan mode showed a significant increase in the Cup-to-disc (C/D) area ratio (+39% at 3Mo and 88% at 5 months post-LPC) and decrease in the Rim area (−57% at 3 months post-LPC and −45% at 5 months post-LPC) most likely reflecting optic nerve damage including both demyelination and axonal loss (Table S6).

### Immunohistological characterization of optic nerve lesions

Next, we analyzed the long-term effect of LPC induced demyelination in the optic nerve by immunohistochemistry and electron microscopy on cross-sections of the LPC-ON and C-ON of the same animals (see Table1) at 3-, 5- and 9 months post-LPC injection.

Immunohistochemical analysis of the optic nerve showed a decrease in the nerve section area of 2, 1.7, 1.9 folds in the LPC-ON over C-ON at 3-, 5- and 9 months post-LPC respectively (Fig. 4A, B, C). Although the global architecture of the LPC-ON appeared to be well preserved, gaps between bundles occurred at the latest time-point (9 months). Immunolabeling for MBP revealed that demyelination was extensive with a decrease in the MBP+ area of 1.3, 1.9, and 2.6 folds in the LPC-ON over C-ON at 3-, 5- and 9 months post-LPC (Fig. 4D, E, F). Immunohistochemistry for NF to detect neurofilaments in axons, showed only partial preservation of axons in the LPC-ON with a decrease in the number of NF+ axons of 1.2, 1.3, and 1.7 folds in the LPC-ON over C-ON at 3-, 5- and 9 months post-LPC (Fig. 4 G, H, I). Double immunodetection of MBP and NF showed that axons were mostly devoid of MBP+ myelin in the core of the lesion, but enwrapped by MBP+ donut-shaped structures, at the lesion edge (Fig 4H), suggesting the presence of re-myelinated axons.

**Fig 4.**
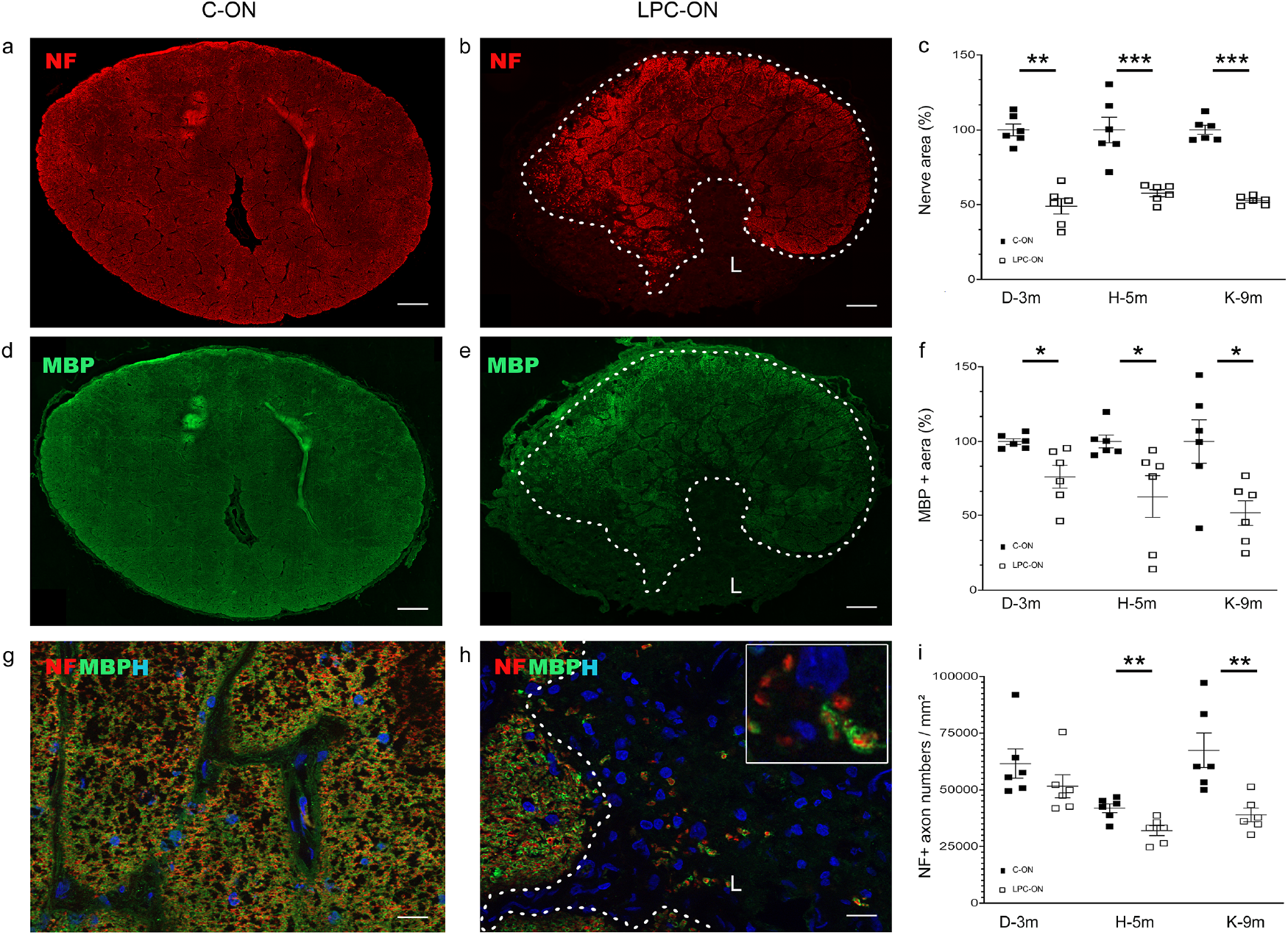
Immunohistological characterization of axons and myelin in 9 months post-LPC C-ON **(A, D, G)** and LPC-ON **(B, E, H)** representative cross-sections. (**A-H**) Immunodetection of axons (NF, red) and myelin (MBP, green) viewed in single- **(A, B, D, E)** or combined- **(G, H)** channels. Dotted lines indicate the lesion (L) limit. **(C, F, I)** Quantifications of the nerve section area (**C**), MBP+ myelin area (**F**), and Neurofilament+ structures **(I)** in C-ON and LPC-ON at 3- (n=1), 5- (n=1,) and 9- (n=1) months post-LPC. Nerve shrinkage correlates with myelin and axonal loss. Inset in (**H)** illustrates the presence of axons enwrapped by MBP+ myelin at the lesion border. For each subject, data for the LPC-ON (empty squares) are expressed In (**C**) and (**F**) as a percentage of the corresponding C-ON fellow nerve values (full squares) and axon number/mm2 in (**I**). Data are the mean ± SEM, t-Test \**P*< 0.05, ** *P*<0.01, \*\*\**P*<0.001. Scale bar **(A, B, D, E)** 200μm, **(G, H)** 20μm. L=lesion.

As remyelination is a complex process involving multiple glial partners, we analyzed the cellular dynamics of the lesion in the C-ON and LPC-ON. Hoechst labeling to detect all cell nuclei, showed a significant increase in cell density of 1.5, 1.3, and 1.4 folds in lesions of the LPC-ON over control, at 3-, 5-, and 9 months post-LPC (Fig. S2A-B’, G). In rodents, OPC (15) and mature oligodendrocytes (29–31) are involved in the process of remyelination. Therefore, we analyzed the oligodendroglial compartment performing double immunolabeling for Olig2 as a general marker of the oligodendroglial lineage and CC1 as a marker of mature oligodendrocytes. While Olig2+ cells were present in all lesions, their number decreased of 5.1, 26.2 and 1.1 folds in lesions of the LPC-ON compared to the C-ON at 3-, 5-, and 9 months post-LPC (Fig. 5A-D, E). This decrease in Olig2+ cells involved immature Olig2+/CC1- and mature Olig2+/CC1+ oligodendroglial cells (Fig. 5 F, G). Immunolabeling for the proliferation marker Ki67 and OLIG2, and Hoechst dye showed that few Hoechst+ or Olig2+ cells were Ki67+ within the lesion including its border at any time-point, suggesting their quiescence. Immunodetection of myeloid cells (CNS-resident microglia and blood-derived monocytes) with the Iba1 marker pointed to a significant increase in Iba1+ cell density of 2, 2.7, 2.7 folds in the LPC-ON over C-ON at 3-, 5- and 9 months post-LPC (Fig. S2H). Interestingly while Iba1+ cells were homogeneously dispersed in the C-ON and LPC-ON, they had features on non-activated microglia with ramified processes in C-ON, but of activated microglial with amoeboid shapes in the LPC-ON (Fig. S2C’, D’). Immunolabeling for GFAP to detect astrocytes highlighted a marked increase of GFAP+ area of 2.8, 2.2, 2.3 folds in the LPC-ON over C-ON at 3-, 5-, and 9 months post-LPC (Fig. S2I). GFAP+ cells within the lesion had enlarged cell processes compared to GFAP+ cells of control-ON, which were stellate with thin ramified processes (Fig. S2E’, F’). Double labeling for GFAP and NF indicated that GFAP reactivity concerned the whole optic nerve section including the lesion, which was largely depleted in axons. These data indicate that both microglial cells and astrocytes contribute to the increased cellularity detected with Hoechst within the lesion.

**Fig 5.**
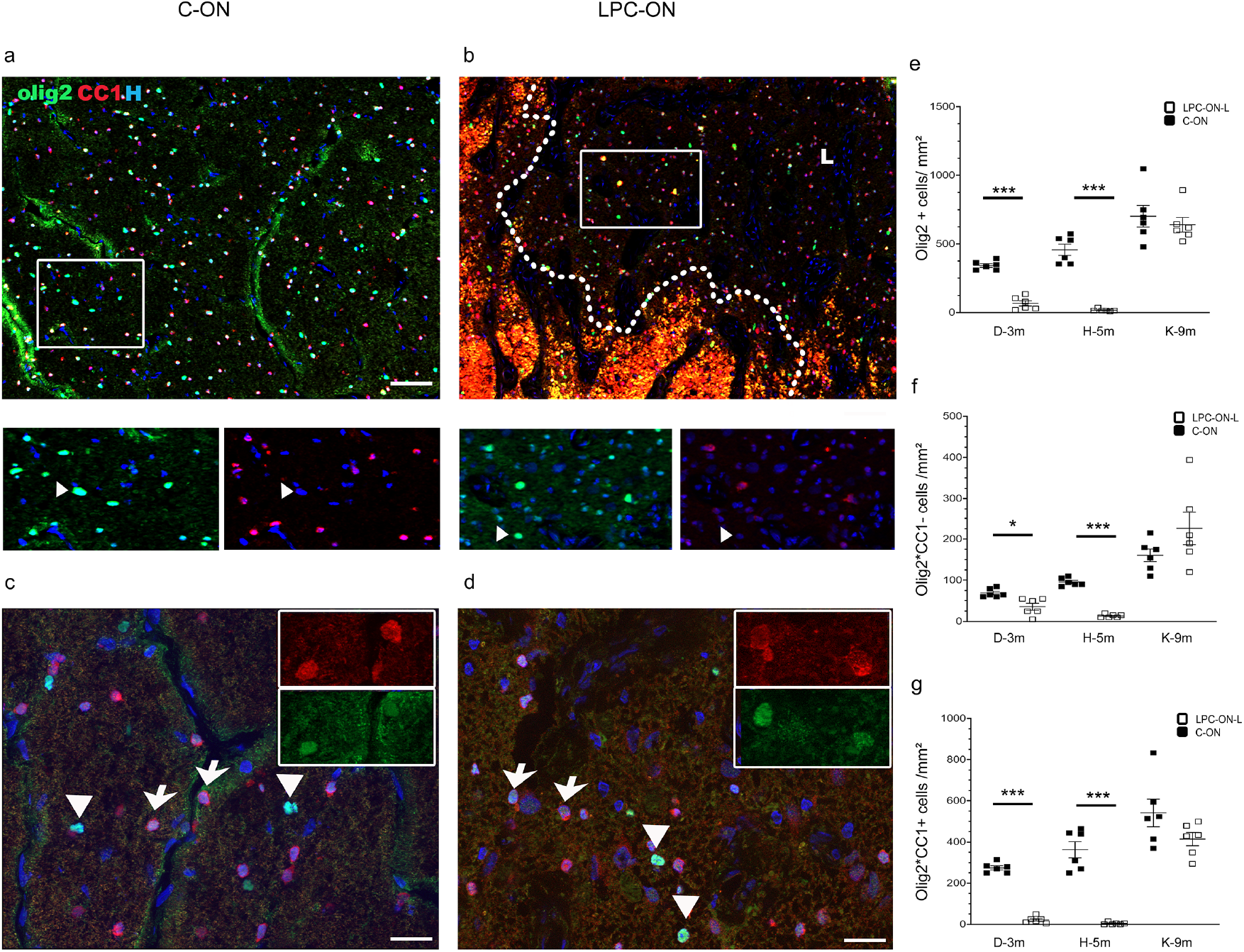
Immunohistological analysis of the optic nerve oligodendroglial populations. Representative 9 months post-LPC cross-sections of C-ON **(A, C)** and LPC-ON **(B, D)** stained for Olig2 (green), CC1 (red), and Hoechst (H, blue). (**A-B**) General views of the parenchyma. Dotted lines indicate the lesion (L) limits. (**C, D**) Details of C-ON **(C)** and LPC-ON (**D**). Insets illustrate single channels. Cells express Olig2 alone (arrowheads) or with CC1 (arrows). (**E, F, G**) Quantifications of Olig2+ (**E**), Olig2+/CC1- (**F**) and Olig2+/CC1+ cells/mm^2^ in C-ON and LPC-ON at 3-, 5- and 9 months post-LPC. Black squares represent values for C-ON and empty squares, values of LPC-ON. Data are the mean ± SEM, t-Test \**P*< 0.05, ** *P*<0.01, *** *P*<0.001. Scale bar **(A, B)** 100μm, (**C, D)** 20μm.

### Ultrastructural changes of the optic nerve lesions

Our previous electron microscopy data showed that demyelination of the optic nerve is achieved at 1week post-LPC injection, leading to healthy demyelinated axon bundles at 1-, 2-, and 6 weeks. While attempts of axon ensheathment by oligodendrocytes were detected at 1- and 2-weeks post-LPC injection, 0.4+0.2% of the surviving axons was surrounded by newly formed myelin (defined by thin myelin sheaths) (28).

Presently, we analyzed the long-term consequence of LPC injection in the optic nerve at 3-, 5-, and 9 months post-LPC. Semi-thin cross-sections stained for toluidine blue confirmed shrinkage of the LPC-ON nerve area, ranging from 26 to 60% of the C-ON area (Fig. 6A-D). At the ultrastructural level, the C-ON harbored myelinated axon bundles separated from each other by vascular structures and perivascular astrocytes, and oligodendrocytes (Fig. 6A1, A2). The LPC-ON was characterized by extensive demyelination. Although scaring or necrosis did not occur, the bundle organization was partially disrupted in the LPC-ON lesion. Axon depletion was more obvious in the lesion core than the rim. While astrocyte processes surrounded axon pockets at 3 months post-lesion (Fig. 6B1), fibrous astrocyte processes progressively occupied the space normally occupied by axons confirming our immunohistological observations (Fig. 6C1-D2, Fig. S3A-F). Moreover, the denuded axons contained neurofilaments, but became depleted in microtubules, a sign of axon sufferance (Fig. 6C1, D1) (41, 42). Electron dense axo-glial contacts were also observed at later time-points (Fig. S3D, F). Myelin-laden macrophages and microglia were present mainly in the lesion rim, (Fig. S4A, D, E). As observed by immunohistochemistry, mature (dark) and immature (light) oligodendrocytes were present at all time points in the lesion core and rim (Fig. S4A, B). In the rim, oligodendrocytes were sometimes in contact with axons without forming myelin. Others were surrounded by remyelinated axons (recognized by thin myelin) (Fig. 6B2, C2, D2). Remyelination was present only at the rim of the lesion. The percentage of remyelinated axons among the surviving axons at the lesion rim was similar among the different time-points (12.66%, 13.39%, 12.18% at 3-, 5-, and 9 months respectively), and only slightly increased compared to 6 weeks post-LPC (25). G ratios of remyelinated axons in the LPC-ON (LPC-ON 3 months: 0.87±0.004, *P*<0.001; 5 months: 0.86±0.006, *P*=0.001; 9 months: 0.86±0.004, P< 0.001; n=6 per nerve/animal) were significantly higher than C-ON (C-ON: 0.73±0.009, n=6 per nerve/animal) and did not seem to vary among animals (*P*: 0.98; 0.90; 0.99, n=6 per nerve/animal). Thus, although LPC injection of the optic nerve leads to demyelination and remyelination onsets at early time points (28), it is followed by major axon loss, persisting inflammation (microglial cells and astrocytosis), and sustained demyelination at later time points.

**Fig 6.**
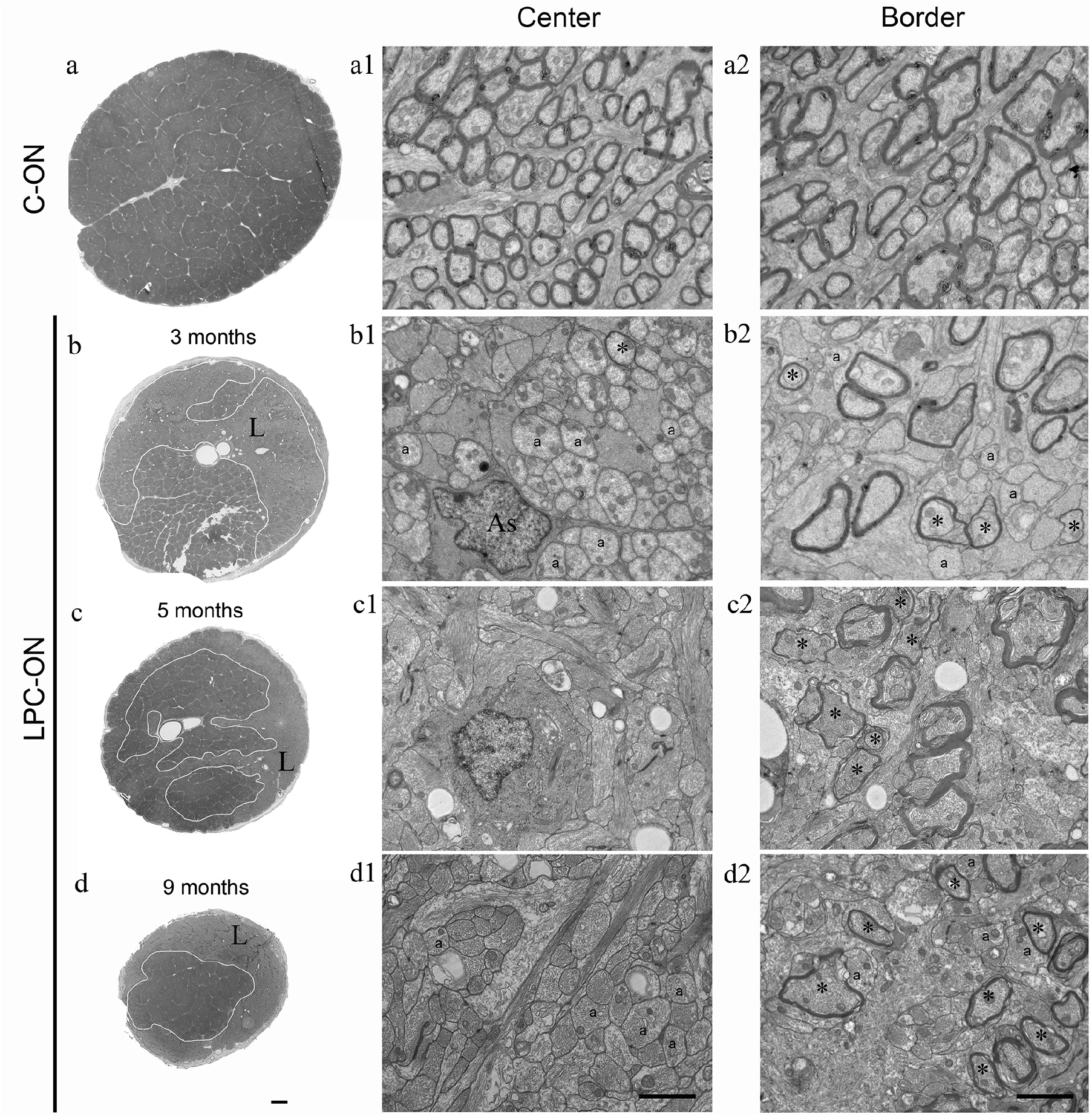
Ultrastructural characterization of the C-ON **(A, A1, A2)** and LPC-ON at 3- **(B, B1, B2)**, 5- **(C, C1, C2)**, and 9- **(D, D1, D2)** months post-LPC. **(A, B, C, D)** Semi-thin optic nerve crosssections illustrating progressive nerve shrinkage. **(B1, C1, D1)** Ultra-thin sections illustrating the center of the lesion. **(B1)** At 3 months post-LPC, a great majority of axons survive and form clusters surrounded by astrocyte (As) processes. **(C1 and D1)**, At 5- and 9 months post-LPC, axon clustering is followed by progressive axon loss and replacement by astrocyte processes. (**B2, C2, D2**) Ultra-thin sections illustrating the border of the lesion. Thin myelin (*) identifies remyelination at the lesion rim only. Scale bar **(A, B, C, D)** 200μm, (A1-D2) 2μm.

### Immunohistological characterization of retinas

As retinal ganglion cells project in the optic nerve, we examined the consequence of long-term ON demyelination on the retina. First, we performed immunohistochemistry on whole-mount retinas (control and lesion side) to generate a global view of the impact of the optic nerve demyelination on the morphological status of the LPC-ON corresponding retina, and compared it to that of the C-ON (Fig.7). Immunolabeling for NF and Brn3a, markers of neurofilaments and RGCs respectively, highlighted a decrease of 2, 3.6, and 1.5 folds in the nerve fiber layer area, as well as a decrease of 5.8, 11.6, and 2.4 folds in RGC density on the LPC-ON side compared to the C-ON side at 3-, 5- and 9 months post-LPC. Analysis of the RGCs versus the RNFL data indicated that RGCs were more vulnerable than RNFL with a decrease of 5.2 folds for RGC and 2.2 folds for RNFL for the LPC-ON compared to C-ON corresponding retinas.

**Fig 7.**
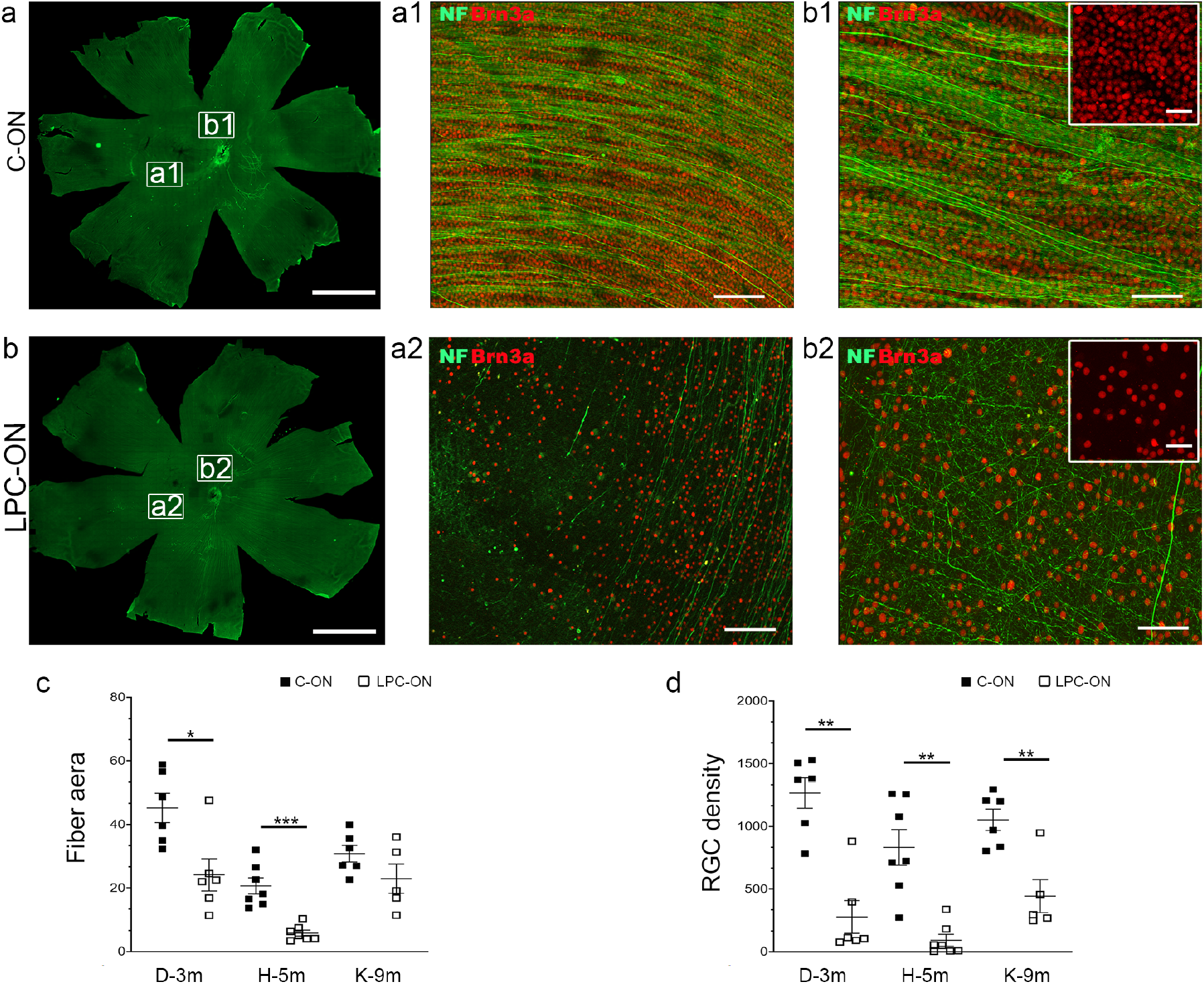
Immunohistochemical characterization of the LPC-induced retinal modifications. (**A, B**) Immunodetection of Brn3a (red) for RGCs, and NF (green) for axons: drastic reduction of RGCs and axons in the LPC-ON corresponding retina **(B, Ba, Bb)**, compared to the C-ON **(A, Aa, Ab)** retina at 9 months post-LPC. Boxed areas near the fovea (**a**) and optic nerve head (**b**) are magnified in **Aa, Ab**, and **Ba Bb** respectively. (**C, D**) Quantification: significant decrease in the nerve fiber area fraction (**C**) and RGC density (**D)** with time post-LPC. Data for each subject are expressed relative to the corresponding C-ON fellow nerve values (n=1 per time-point). Data are the mean ± SEM, t-Test *** *P*<0.001. Scale bar **(A, B)** 6mm, **(Aa, Ab, Ba, Bb)** 50μm, (inset) 25μm.

### Ultrastructural retinal changes

Alterations of the retina were further validated in semi-thin and ultra-thin cross-sections of two animals sacrificed at 5- and 9 months post-LPC. Semi-thin crosssections of the retina revealed a progressive decrease in retinal thickness post-LPC (C-ON: 310μm; 5 months: 260μm; 9 months: 200μm) involving the inner most retinal layers of optic nerve fibers (layer 9) and RGC (layer 8) (Fig. 8). Analysis of ultra-thin sections confirmed that thickness reduction concerned the retinal inner layers compartment (C-ON: 150μm; LPC-ON 5 months: 105μm; 9 months: 60μm) including the nerve fiber-, and ganglion cell-layers, while the outer layer thickness remained unaffected (C-ON: 160μm; LPC-ON: 5 months: 155μm; 9 months: 140μm) (Fig. S5). Anomalies of the retinal inner layer were further characterized at the ultrastructural level. The nerve fiber-, retinal ganglion cell-, and inner plexiform layers all showed various degrees of alterations. Axon bundles were severely disorganized in the retinal fiber layer, 5 months post-LPC, and completely vanished, 9 months post-LPC. The major loss occurred in the RGC layer at 9 months post-LPC compared to 5 months post-LPC. Moreover, a decrease in synaptic contacts and altered mitochondria in the synaptic endings occurred in the plexiform layer and was more severe at 9 months post-LPC compared to 5 months post-LPC. By contrast, the retinal pigmented epithelium, photoreceptor layer, outer limiting membrane, and outer plexiform layer had a very similar morphology in the LPC-ON corresponding retinas compared to those of the C-ON. These morphological observations indicate that the LPC-induced chronic lesions of the optic nerve were associated with progressive retinal degeneration of the inner retinal layers.

**Fig 8.**
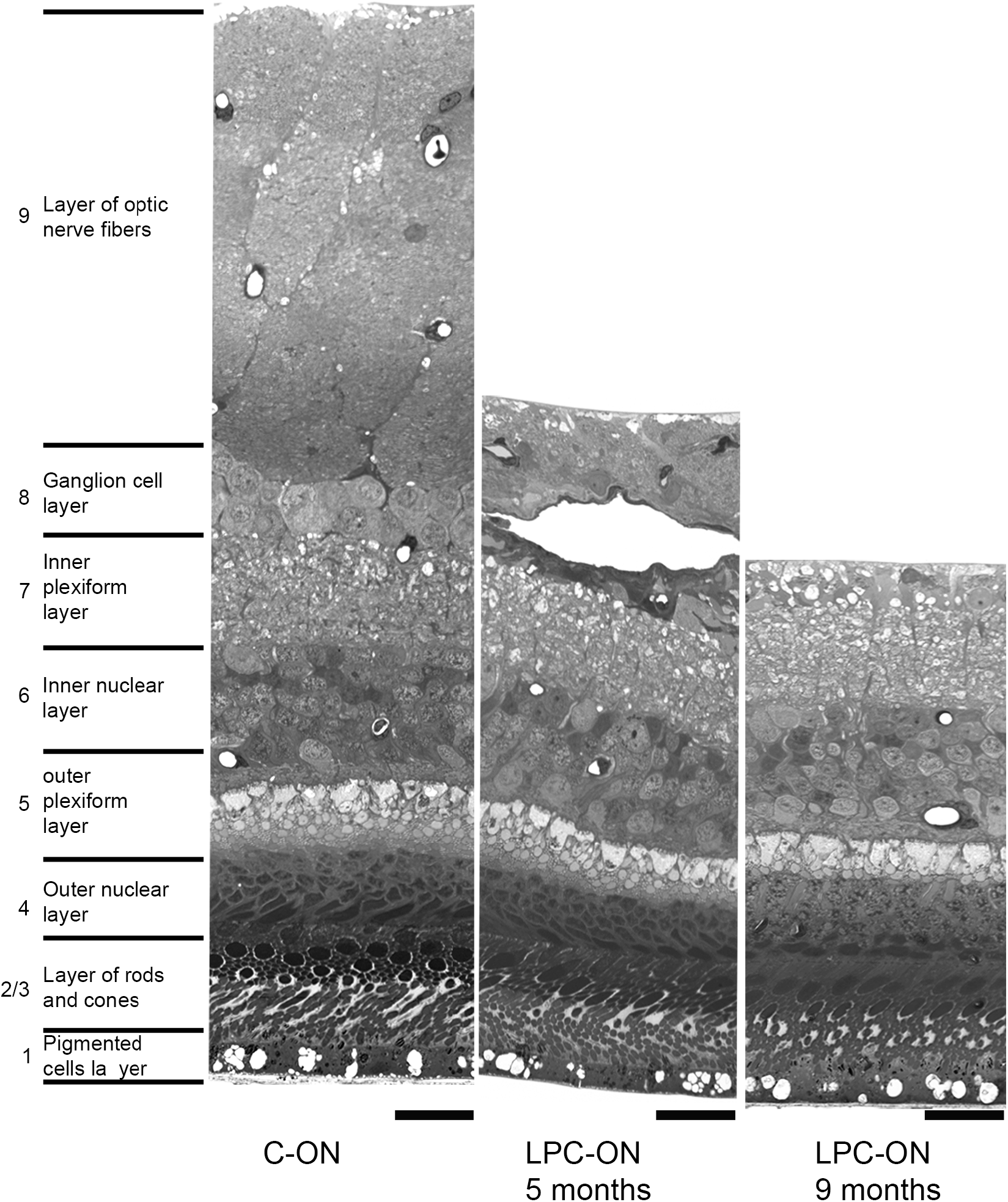
Analysis of the LPC-induced retinal modifications on semi-thin cross-sections: progressive decrease of the retinal layers 8 and 9, at 5- and 9 months post-LPC compared to the C-ON retinal layers. Scale bar: 25μm.

## Discussion

Since we initiated this work, several pre-clinical studies addressed the consequences of optic nerve myelin perturbations on visual functions in rodents, dogs, and cats (18–23). Yet the relevance of these correlative studies to primates is questionable in view of several distinct differences existing between these species and primates (12). In particular, old-world monkeys have eye sizes much closer to man, visual acuity driven macula which is absent in mice and reduced in dogs, and are endowed with trichromatic vision similar to humans. Another advantage of non-human primates is the availability of a panel of instruments that can reveal insights into the visual system state in living rather than anesthetized conditions. VF, VEP, and PLR on awake animals can be correlated to structural assays such as OCT and ERG, all of which are used in clinic for altered vision disorders including MS. For example, while compression or genetically acquired hypomyelination of the optic nerve has been associated with altered VEPs in mice 18-21) and dogs (23), these assays consisted in flash recordings and were always performed on anesthetized subjects, a condition likely to modify VEP and ERG responses (32). Moreover, none of these pre-clinical studies addressed the functional consequences of long-term optic nerve demyelination in non-human primates. Combining these approaches, we show that LPC-induced demyelination of the macaque optic nerve leads to altered VF, VEP and PLR responses but preserved ERG. PLR and VEP were predictive of optic nerve lesions involving the RGC population, and this was confirmed both by OCT and post-mortem analysis. While VEP assays address the visual pathway, PLR, relates to the non-visual pathway, which is mediated by subset of RGC. Our data suggest that the optic nerve lesions affected both pathways. ERG predicted the absence of photoreceptor and bipolar cell involvement and thus the absence of retrograde degeneration, all of which was confirmed by post-mortem analysis as well.

In clinical studies involving MS subjects, it is uncertain whether VEP alterations as a mean to assess axon conduction are correlated anatomically to optic nerve demyelination and/or axonal loss (33). We show that the delayed VEP responses correlated with both, optic nerve demyelination and axonal loss at the histological level. In our earlier study, we showed that demyelination of the optic nerve occurs as early as 1week post-LPC injection and is associated with poor remyelination in spite of fair axon preservation up to 2 months post-LPC injection (28). Using the same protocol and operator, we presently show by immunohistochemistry and electron microscopy, that remyelination stalls and axonal loss worsens from 3 months post-LPC (Fig. S6). Thus the early phase of demyelination/remyelination with minor axonal loss is followed by a second phase of chronic demyelination and axon degeneration. Interestingly, delayed VEP responses appeared as early as 1month post-LPC, a time at which the majority of axons were demyelinated and appeared healthy, suggesting that the early VEP response is caused by demyelination rather than axonal loss. We also show at the histological level that the second phase (5 and 9 months post-LPC) is characterized by the progressive replacement of axons by astrocyte processes and persisting demyelination. Interestingly, VEP latency restoration or aggravation overtime was not observed. While the absence of VEP restoration is consistent with the absence of axon regeneration and remyelination, the absence of VEP worsening could be accounted for environmental modifications such as astrogliosis and glial-axon densities that were prominent in the LPC-ON and corresponding retina, at later time-points. These increased membrane relationships between axons and glia are characteristics of persisting demyelination in chronic EAE (optic nerve and spinal cord) and MS plaques (spinal cord) (34, 35), and may confer a supportive role of glial cells to axons, thus representing a compensatory mechanism to the impaired nerve fiber conduction (34, 36).

As RGC project into the optic nerve, we questioned the consequence of optic nerve demyelination on the retina. SD-OCT on 2 subjects highlighted thinning of the RNFL at 3 and 5 months with no significant differences between the 2 time-points, and retinal thinning was further validated by a decrease in RGC densities and retinal axons at the immuno-histological level, and thinning of the inner retinal layers at the ultrastructural level. SD-OCT is a non-invasive method of choice to track neuronal-axonal loss, and has provided evidence for injury to the inner retinal layers including the RNFL and reduction of RGC cell density in MS patients (37). Our model thus validates that RNFL thinning reflects both neuronal and axonal loss. Retinal thinning occurs in MS with- (38) or without optic neuritis (39), and as early as 6 months after the first optic neuritis episode (38). The time-lines of retinal damages in response to the macaque optic nerve demyelination model is nearly comparable to man when adjusting macaque’s mean life-duration to man. Interestingly, RNFL thinning was not observed in conditions of optic nerve hypomyelination in dogs (23), raising the question of RNFL thinning being a primary or secondary event to optic nerve demyelination. Our data showing that a primary event of optic nerve demyelination leading to optic nerve axonal loss is followed by RNFL thinning as seen by OCT and histological evaluation, corroborates the hypothesis that an optic nerve insult can be the cause of retinal pathology.

ERG predicted sparing of the outer retinal layers, a finding which was also confirmed at the immuno-histological and ultrastructural levels, thus indicating that the optic nerve induced demyelination has spared rods and cones. ERGs are often affected in MS (40) and could result either from retrograde trans-synaptic degeneration of the retinal outer layers secondary to optic nerve atrophy or for a small patient subset, retinal auto-immunity as a possible mechanism of degeneration (7). Although in the non-human primate model, ERGs were normal and sparing of the retinal outer layer was validated by electron microscopy, the possibility that trans-synaptic degeneration of the outer retinal layers leading to altered ERG, could occur at later times cannot be excluded.

Although remyelination onsets during the first month post demyelination in the nonhuman primate optic nerve (28) our present study indicates that it is compromised over time. Reasons for this failure could be multiple. While in rodents, remyelination is achieved successfully after acute demyelination even in the optic nerve, it relies on oligodendrocyte progenitor proliferation, recruitment and differentiation into mature oligodendrocytes (41). This initial burst in oligodendrocytes did not occur in the present model (28) sustaining its similarity with MS chronic lesions (42–44) and experimental models of chronic demyelination (45, 46). Although depleted in numbers, mature and immature oligodendrocytes were seen within lesions up to 9 months post-demyelination. Both the slow tempo of differentiation of primate oligodendroglial cells (47–49) and their lack of proliferation, possibly caused by delayed astrocyte recruitment (28) could have prevented oligodendrocyte timely differentiation into myelin-forming cells. Moreover, noxious inflammatory conditions sustained by myeloïd cells (monocytes and microglia), known to modulate remyelination (15), could also have blocked oligodendrocyte maturation as demonstrated for rodent and human oligodendrocytes (50–53). While at early stages, macrophages/microglial cells were widely spread within the optic nerve lesions, they became restricted to the lesion rim at later stages, as observed in chronic or inactive MS lesions (54). Although present at early times, they may have failed to switch timely from a pro- to an anti-inflammatory phenotype to allow remyelination to proceed (55,56).

Failed remyelination was associated with progressive axonal loss and retinal ganglion cell degeneration. Although axons, were present in fair amounts and appeared healthy up to 2 months post-demyelination (28, Fig. S6), their number decreased severely from 3 months on (present study) with no significant differences among animals and correlating with optic nerve shrinkage. At the ultrastructural level, surviving axons remained anatomically normal up to 3 months but were characterized thereafter (5 months) by a number of cytoplasmic anomalies, including increased axonal organelles, loss of neurofilament spacing and microtubules as described for chronic MS lesions (57). While the cause of axonal loss and degeneration and their precise timing remain to be further investigated, sustained inadequate inflammation, glutamate excitotoxicity and reduced oligodendroglial trophic support to axons (1, 57), may all have contributed to the progressive axon degeneration and evolution of the lesion from an early active state with remyelination onset, to a late chronic state with failed remyelination, neurodegeneration and neurological dysfunction, mimicking the many features of MS chronic lesions (1, 57–59).

### Conclusion

Although this non-human primate model of optic nerve demyelination does not represent the auto-immune component of MS, or its correlated pre-clinical model EAE, it has the obvious advantage to target the optic nerve in a very predictive manner with demyelination followed by the onset of remyelination at early time points and failed remyelination, and axonal loss and visual dysfunction at late time-points. The present model could provide a platform to improve our understanding of the mechanisms underlying axon pathology progression and failed remyelination. Moreover, the existence of a correlation between the functional assays and the structural/anatomical observations makes this model an excellent tool to develop strategies aiming at promoting neuroprotection and/or remyelination, to overcome the inevitable neurological disabilities of myelin diseases such as MS.

## Materials and Methods

### Animals

Adult male *macaca fascicularis* monkeys (5-10 years old) were purchased from and raised by Silabe, Strasbourg or Bioprim, Perpignan, France, and hosted for the time of the experiments at the ICM primate facility. For ethical reasons, the number of animals (n=6) was kept as minimal as possible following the legal requirements. The identification codes refer to individual monkeys (Table S1). During quantification, animals had another code preventing the experimenter to identify the animals. Experiments were performed according to European Community regulations and INSERM Ethical Committee (authorization 75-348; 20/04/2005) and approved by the local Darwin ethical and National committees (authorization # 3469, and # 17101).

### Demyelination

LPC injection was performed in one optic nerve, the other optic nerve serving as control. Saline injection was reported to have minimal impact on demyelination and axon loss compared to LPC injection (28). Details can be found in *SI Methods*.

### Electroretinogram

ERG recordings were performed under anesthesia (with a mixture of ketamine (3mg/kg)/Dexdomitor (0,015mg/kg) in two occasions, once before surgery (baseline) and once after, and are the mean of 3-4 animals. The method for ERG recordings (60) is described in detail in *SI Methods*.

### Pupillary light reflex

Changes in PLR parameters, such as constriction amplitude and latency, have been associated with retinal dysfunction, including degeneration of the rod and cone photoreceptors (61), and melanopsin-containing ganglion cells (62). The constriction and dilatation of the pupil in response to changes in light intensity were evoked and collected on awake animals by a portable SIEM-BIOMEDICAL device (Biovision) equipped with an infrared-sensitive camera delivering continuous infrared illumination generating pupil images at 60 frames/sec. Details are provided in *SI Methods*.

### Visual evoked potential

VEP recordings (63, 64) were performed in a dark room, using the ©MetroVision system, on awake animals using checkerboard and flash stimulations (750 msec) delivered bi- and monocularly averaging 60 sweeps per trial. Stimuli were 1 to 2 Hz. D15’ and D30’ checkerboard- and red and blue flash responses were selected over D60’ and grey flash respectively, as the most sensitive and reliable parameters of follow up throughout the study. Details can be found in *SI Methods*

### Visual field

VF sensitivity was indirectly assessed before and after the lesion using saccadic responses to targets presented successively in the visual field. This procedure was used in monocular and binocular conditions with awake monkeys placed at 60 cm from a CRT screen, trained and rewarded to make saccades to visual targets using the Jeda in-house software (65). Details are provided in *SI Methods*.

### Optical coherence tomography

OCT imaging was performed on a spectral-domain iVue (software version 3.2; Optovue, Fremont, CA).and was conducted under anesthesia in 2 animals after appropriate dilatation by Tropicamide 0.5% (Mydriaticum, FARMILA-THEA FARMACEUTICI, Italy). The detailed method can be found in *SI Methods*.

### Tissue processing

Animals were sacrificed by deep anesthesia at 3- (n=2), 5- (n=2), and 9- (n=2) months after LPC injection in the left optic nerve. The right optic nerve and retina were used as internal controls. Details are provided in *SI Methods*.

### Immunohistochemical and ultrastructural quantifications

Quantifications were performed as described in details in *SI Methods*.

### Statistics

were performed as descibed in *SI Methods*.

## Acknowledgments

We thank Morgane Weissenburger, Marion Lanoe, Harry Ahnine, Oceane Aribo, and Serban Morosan from the ICM Primate Facility, Dominique Langui, from the ICM Imaging Facility as well as Fabrice Arcizet and Celine Nouvel-Jaillard from IDV for their valuable operational contributions and support. We are grateful to Dr. A. Green and members of his team for helpful discussions, and Dr S. Mozafari for her valuable advice on remyelination assessment. This work was supported by the programs ‘Institut des Neurosciences Translationnelles’ ANR-10-IAIHU-06, the International Associated Laboratory “Neuro-Bridge” and the NIH/NINDS SRO1NS105741-02.

## Author Contributions

NS coordinated, designed, and performed most of the experimental work, analyzed, interpreted the results, and wrote the manuscript. PM performed the optic nerve surgery. ECR and SG, performed the animal task training. SR (ERG), EB (OCT), ECR (PLR) performed and interpreted the functional assays. MF performed the immunohistochemical imaging and statistical analysis. CB performed the electron microscopy processing, imaging and analysis. RA and PP carried out the functional assay MATLAB analysis. LYC performed the functional study statistical analysis. PP, SR, SP, JL, advised on the functional assays and critically reviewed the manuscript; ABVE conceived the project, handled the funding, supervised and interpreted the results, wrote and edited the manuscript with critical input from all authors

## Supplementary Information

### SI Methods

#### Demyelination

Prior anaesthesia each animal received an intra-muscular injection of antibiotic (Amoxicilline, 15mg/kg, Zoetis) followed by an analgesic (buprenorphine (0.03mg/kg, Axience). Anaesthesia was induced with a mixture of ketamine (3mg/kg, VirBac)/Dexdomitor (0,015mg/kg, Vetquinol) and animals were maintained under Isofluorane 1.5-2% during surgery. The temperature was maintained constant during the whole procedure. The cardiac and respiratory functions were monitored as well. Access to the optic nerve was achieved by a lateral approach of the eye orbit. The left optic nerve was exposed by resection of the lateral orbital bone, retraction between rectus superior and rectus lateralis muscles, peri-ocular fat removal, and incision of the dura. Demyelination was induced by injecting 2 × 1μl of a solution of 1% LPC (Sigma, Saint Quentin Fallavier, France) at a speed of 0.5 μl/min via a glass micro-pipet (300-μm diameter) connected to a Hamilton syringe. The gliotoxin was injected 5 mm caudally to the *lamina cribrosa* in the optic nerve.

#### Electroretinogram

ERG recordings were performed under anesthesia (with a mixture of ketamine (3mg/kg) /Dexdomitor (0,015mg/kg) in two occasions, once before surgery(baseline) and once after, and are the mean of 3-4 animals. Animals were placed in a dim light room, their head immobilized with the use of a restrainer. The eyes were maintained open with the use of a sclero-conjunctival atraumatic needle, which also served as active ERG electrode. Drops of tropicamide (Mydriaticum ^ND^ Théa, France) were applied to each eye to obtain a stable mydriasis. Corneal hydration was maintained throughout the entire procedure with the use of carbachol (Ocrygel ^ND^ TVM, France). A *Visiosystem* (10.2.57, Siem Bio-Médicale, Nîmes, France) was used to generate the flash stimuli, as well as to record and analyze the ERG responses. Binocular full-field ERGs were elicited with 2 photo-stimulators (source: achromatic LEDs) for background conditions and flash stimulations (maximum intensity: 1.9 log cds/m^2^) in both photopic and scotopic conditions. The cone system was tested in photopic conditions against a bright background (25 cd.m^2^) aimed at desensitizing the rod system during 10 minutes. The flash ERG responses were obtained with 9 decreasing intensities of a series of 15 white LED bright flashes stimuli (from 1.90 log cd.s.m^2^ to 0.80 log cd.s.m^2^) delivered at 1.3 Hz temporal frequency (inter-stimuli interval of 769 ms) to determine the “Imax” intensity corresponding to the maximum b-wave amplitude (V max) observed at the saturation point of the luminance curve (maximum cone system response). The flicker ERG responses were obtained with the white flash stimulus at Imax intensity as described above and delivered at 30 Hz temporal frequency during at least 15 seconds. Then, the light was switched off and the rod system was tested in scotopic conditions. After 20 minutes of dark adaptation, flash scotopic responses were obtained in the dark with an average of 5 dim light flashes (intensity: −1.1 log cd.s.m^2^) delivered at 0.1 Hz temporal frequency, corresponding to 10-second inter-stimuli intervals. Two minutes after the last scotopic flash, in the same scotopic conditions, the combined rod-cone response was elicited with a unique white flash at Imax intensity as described above.

For the photopic response analysis (photopic stimuli, photopic background), a-wave and b-wave amplitudes and implicit times (peak times) were measured in order to determine the maximal cone responses (V max at I max). Values were considered as representative of cones system function for flash and flicker responses. The amplitude values depend on the background intensity: when the background intensity increases, the amplitude values decrease.

For the scotopic response analysis (scotopic stimulus, scotopic background), b-wave amplitudes and implicit times (peak times) were measured and considered as representative of rods system function. The values measured at 20 minutes of dark-adaptation were considered as representative of rods-dominated system function.

For the combined responses (photopic stimulus using I max, scotopic background), a-wave and b-wave amplitudes and implicit times (peak times) were measured, and values obtained using I max intensity were considered as representative of combined (mixed) rod-cone responses.

#### Pupillary light reflex

The direct PLR was evaluated for each eye and each condition using a series of 1-sec. illuminations elicited with increasing blue-light (BL) (480 nm) and decreasing red-light (RL) (630 nm) intensities (n=7, with minimum: −0.02 log cd/m^2^ to maximum: 2.96 log cd/m^2^). PLR responses were recorded 1 sec before stimulation to establish the baseline, and for 9 sec. after stimulation, and included the following parameters: PLR constriction amplitude (C2 peak), Post-Illumination Pupil Response (PIPR), PLR latency, and PLR constriction time. Amplitude measurements were expressed as a percentage of the baseline pupil surface (pixel). PIPR was deduced from differences between the generated curve and baseline. The C2 peak and PIPR data were analyzed using a routine Matlab custom software. For each time point, data represent the mean values ± SEM recorded from both electrodes over 2 weeks before, and 2 weeks after the date, at 1-, 3-, 5-, and 9 months post-lesion.

#### Visual field

Once monkeys fixed a target located in the screen center, a new randomly chosen target, was presented at one of the 8 eccentric locations in a back-and-forth mode. Light stimuli were 0.6021 Cd/m^2^, 3.5726 Cd/m^2^, 8.6250 Cd/m^2^. Monkeys made saccades to these back-and-forth successions of targets, in order to probe the entire visual field. As unseen targets entail long saccadic latencies, latencies (msec) were analyzed and used to derive the monkey’s sensitivity before and after the lesion. Data were analyzed using a routine Matlab custom software, and represent the mean values ± SEM, between the minimal and maximal values excluding values > 15 sec., over 2 weeks before and 2 weeks after the chosen date of 1-, 3-, 5-, and 9 months postlesion.

#### Visual evoked potential

Two active electroencephalogram (EEG) type electrodes were placed on the scalp, at Oz which is the highest point of the occiput, laying over the visual cortex. The reference and ground electrodes were placed at Fz and Cz, respectively. Black and white checkerboards (15’, 30’, 60’) were followed by achromatic flashes (red, blue, and gray). Stimuli were 1 to 2 Hz. Checkerboard parameters included: N50, P80, and N130 waves. Flash parameters included: N1, P1, N2, P2, N,3, and P.

Peak time latencies (msec) were analyzed using a routine Matlab custom software. For checkerboards, the maximal positive peak (P80) time window was defined between 30-137 msec. For flashes, the maximal negative peak (N2) time window 30-120 msec for flashes. For each time point, data represent the mean of values recorded by both electrodes over 2 weeks before and 2 weeks after the date, at 1-, 3-, 5-, and 9 months post-lesion.

#### Optical coherence tomography

Anaesthesia was induced with ketamine (3mg/kg)/Dexdomitor (0,015mg/kg) and animals were maintained under isoflurane 1.5-2% during the procedure. The OCT device *for optical coherence tomography scanning* uses a laser diode with 840 ± 10nm wavelength captures 26.000 A-scans/second with an axial resolution of 5μm. Images with scan quality index (SQI) less than 40 were excluded from further analysis. The Optic Disc Metrics, retinal nerve fiber layer (RNFL) Mapping with Change, Symmetry, and Normative Comparison Analyses were conducted. The average peripapillary RNFL thickness and optic nerve head (ONH) parameters were assessed using the Glaucoma scan mode. It consists of 12 radial scans of 3.4 mm in length (459 A-scans each), and 13 concentric ring scans ranging from 1.3 to 4.3 mm in diameter (429 to 969 A-scans each). All measures were centered on the optic disc.

The *Retina map scan pattern* was used for the retinal thickness measurements. A 6 mm^2^ area of the retina was scanned with horizontal scan lines of 250-micron separation, each consisting of 512 A-scans per line. The Retina map mode includes the high-definition scan pattern “Hi-res 7-line raster” consisting of 1024 A-scans, 6 mm in length, localized in the middle of the scanned area. In this mode the retinal thickness is automatically calculated in the 9 ETDRS-type areas, consisting of a central circular zone 1-mm in diameter and inner and outer rings of 3 and 5 mm in diameter, respectively. The inner and the outer rings are divided into four quadrants: superior, nasal, inferior, and temporal.

#### Tissue processing

*For immunohistochemistry*, animals were perfused with 1L/kg of phosphate-buffered saline (PBS) containing heparin (3ml, 5000 U.I/ml) followed by the same volume of a solution of 4% paraformaldehyde (PFA, Merck, Darmstadt, Germany) in 0.1M PBS. Optic nerves were post-fixed for 24 hours in the same fixative and then soaked for 4 hours in a 20% sucrose solution in 0.1M PBS. Tissues were embedded in OCT medium (Tissue Tek, Sakura Finetek, Torrance, CA), frozen in melting isopentane (−70°C), and cut at 12μm with a Leica Microsystem cryostat. For immuno-labeling, frozen sections were rehydrated in 0.1M PBS. They were then incubated with the primary antibodies diluted in PBS 0.1M supplemented with 4% bovine serum albumin (BSA) overnight at 4°C, washed, and incubated for two hours at RT with the appropriate secondary antibodies. After several washes in 0.1M PBS, sections were counterstained with Hoechst, mounted in fluoromount (Southern Biotechnology, Birmingham, Ala). Primary antibodies were as follows: MBP Rabbit (Millipore, AB980) 1/200 to detect myelin, Olig2 rabbit (Millipore, AB9610) 1/200 for oligodendroglial cells, CC1 mouse (Calbiochem, OP808) 1/200 for mature oligodendrocytes, GFAP: Rabbit (Dako, Z0334) 1/1000 for astrocytes, NF (2F11) mouse (Dako, MO762) 1/200 for axons, SMI31 mouse (Calbiochem, NE1023) 1/200 and SMI32: mouse (Calbiochem, NE1022) 1/200 for phosphorylated and non-phosphorylated neurofilaments respectively, and Iba1 rabbit (Dako, 019-19741) 1/200 for macrophages. For quantification, section scanning, cell visualization, and imaging were performed with the automated AxioScanZ1. For illustration, some images were acquired with a Carl Zeiss Apotome microscope. Retinas were dissected, post-fixed in PFA 4% at 4°C for 24 hours and cleared from the sclera. After rinsing in PBS, whole retinas were immunolabeled overnight at 4°C, with anti-2F11 to label axons (mouse, Dako, MO762, 1/100) and anti-Brn3a (goat, Santa Cruz Biotechnology, sc-31984, 1/750), to label the majority (>75%) of the retinal ganglion cells (RGC) population (1), and incubated with appropriate secondary antibodies. Retinas were then counterstained with Hoechst, flat-mounted, and scanned as above.

*For electron microscopy*, animals were sacrificed by deep anesthesia as above at 3- and 9 months post-LPC injection and were perfused intra-cardially with 1L/kg of phosphate-buffered 0.1M (PB) containing heparin (3ml, 5000 U.I/ml), followed by the same volume of a solution containing 5 % glutaraldehyde, 4% PFA (Sigma) in PB. Tissues (eyes and optic nerves) were removed and immersed for 4 hours in the same solution. Optic nerves were cut transversally into 700-μm thick serial slices. Slices were post-fixed in 2% osmium tetroxide (Euromedex, France) and dehydrated in graded alcohols prior embedding in Epon. One-micron semi-thin sections were stained with toluidine blue and examined in a DMRD Leica microscope. Ultra-thin sections (70 nm) were contrasted with lead citrate and examined with a transmission electron microscope (TEM)-HITACHI 120kV. Retinas were post-fixed in 2.5% glutaraldehyde in 0.1M PB for 24 hours and fixed in 2% osmium tetroxide for 1h at room temperature. Tissues were dehydrated in graded alcohols and embedded in Epon between two silicone slides. Four small pieces of the retina, randomly distributed around the optic nerf entry, were excised and cut transversally in semi-thin (0.5 μm) and ultra-thin (70 nm) sections. Semi-thin sections were stained with toluidine blue for light microscopy. Ultra-thin sections were contrasted with lead citrate for TEM.

The various cell types were identified by their ultrastructural features in the optic nerve and retinal layers. Astrocytes were defined as light cells in appearance with the bean-shaped or irregular nucleus, intermediate filaments (8-9 nm of diameter), glycogen granules, few microtubules, and heterogeneously lined out processes. Oligodendrocytes were distinguished from astrocytes by the greater density of their cytoplasm, round-shaped nucleus, absence of intermediate filaments and glycogen in their cytoplasm, and presence of microtubules (25 nm) in their processes. Macrophages were defined as cells darker than oligodendrocytes, without filaments but with rare microtubules, glycogen, narrow granular endoplasmic reticulum cisternae, the nucleus of variable shapes, and irregular membrane contours or finger-like processes. Their cytoplasm often contains lipid droplets or myelin debris. Axons were defined by the presence of microtubules, intermediate filaments, mitochondria, and regular cross-section profiles.

#### Histological quantifications

*For immunohistochemistry, optic nerve section, lesion areas, and glial populations* were evaluated on immuno-labeled tissue sections, and data for each item collected from 2 scanned sections at 3 levels of the optic nerve lesion (n=6 tissue sections/optic nerve). Image processing was performed with Fiji (2), open-source software for biological-image analysis (36) using in-house-made macros. Lesion areas were defined based upon Hoechst+ nuclei high density and MBP depletion. First, images were binarized, holes were filled and contours defined for precise surface measurements of the whole nerve section (healthy and lesioned nerve) as well as the lesion area. The same contours were then used to quantify the other markers. Myelin (MBP) and neurofilaments (NF) were quantified as a surface fraction using an adapted threshold comparing with the original image. Cell density was automatically calculated by binarization of Hoechst images, followed by segmentation of close nucleus and detection of positive structures with a pre-selected size range. Microglial cells were counted by background subtraction, followed by cell signal optimization before binarization, and segmentation. Hoechst was used as a mask to eliminate false positive microglial cells. For axon and oligodendrocyte counts, backgrounds were established automatically, then areas and thresholds were defined manually. The number of NF+ or Olig2/CC1 positive cells were evaluated over the surface area of C-ON, and LPC-ON respectively. All data are expressed as the mean ± SEM.

##### Retinal nerve layer and ganglion cell population

The density of immuno-labeled RGC population and nerve fiber layer were evaluated on flat-mounted retina scanned at 10X using the Apotome 3 mode (Z stacks, n=10/ RGC and nerve layers respectively). Nine images of 7 areas (n=21) around the optic nerve and macula areas constituting the central retina were randomly selected for quantification of Brn3a+ cells and NF+ areas using the computer-assisted ImageJ program. RGC+ cell numbers and NF positivity were normalized to the surface area. Data are expressed as the mean +/- SEM for each animal.

##### Remyelination

The percentage of remyelinated axons over viable axons and g ratios estimations (myelin diameters over myelin + axon diameters) were performed on ultrathin sections at the middle level of the lesion, and at the lesion rim. The percentage of remyelinated axons was evaluated for axons larger than 1μm diameter. G ratio measurements were performed with ImageJ for at least 80-100 axons per animal. Images were taken from 10-12 randomly selected fields per animal as previously described (3). Data are presented as the mean ± SEM for each animal.

#### Statistics

*For VF, VEP and PLR*, statistical analysis was performed using R (R Core Development Team, 2020, using repeated measures linear mixed-effects models (lme4 package) with “time” (baseline versus follow-up) and “eye” as a within-subjects factor and “time:eye” interaction term for fixed effects. A subject random intercept was included to control for within-subject variability. Parametric tests (ANOVA) were subsequently used to investigate differences between “eyes” at each time point. *For OCT and Immunohistochemistry*. Paired t-Test, bilateral distribution, for small samples was used for two-group comparisons. For immunohistochemistry, it was applied to 5-7 sections analyzed for each monkey as previously reported for non-human primate assessment of axon regeneration (4, 5). Ordinary One-way Anova-Tukey’s multiple comparisons was applied for remyelination assessment. For all assays, *P*-value < 0.05 was set as significance level.

**Fig. S1.**
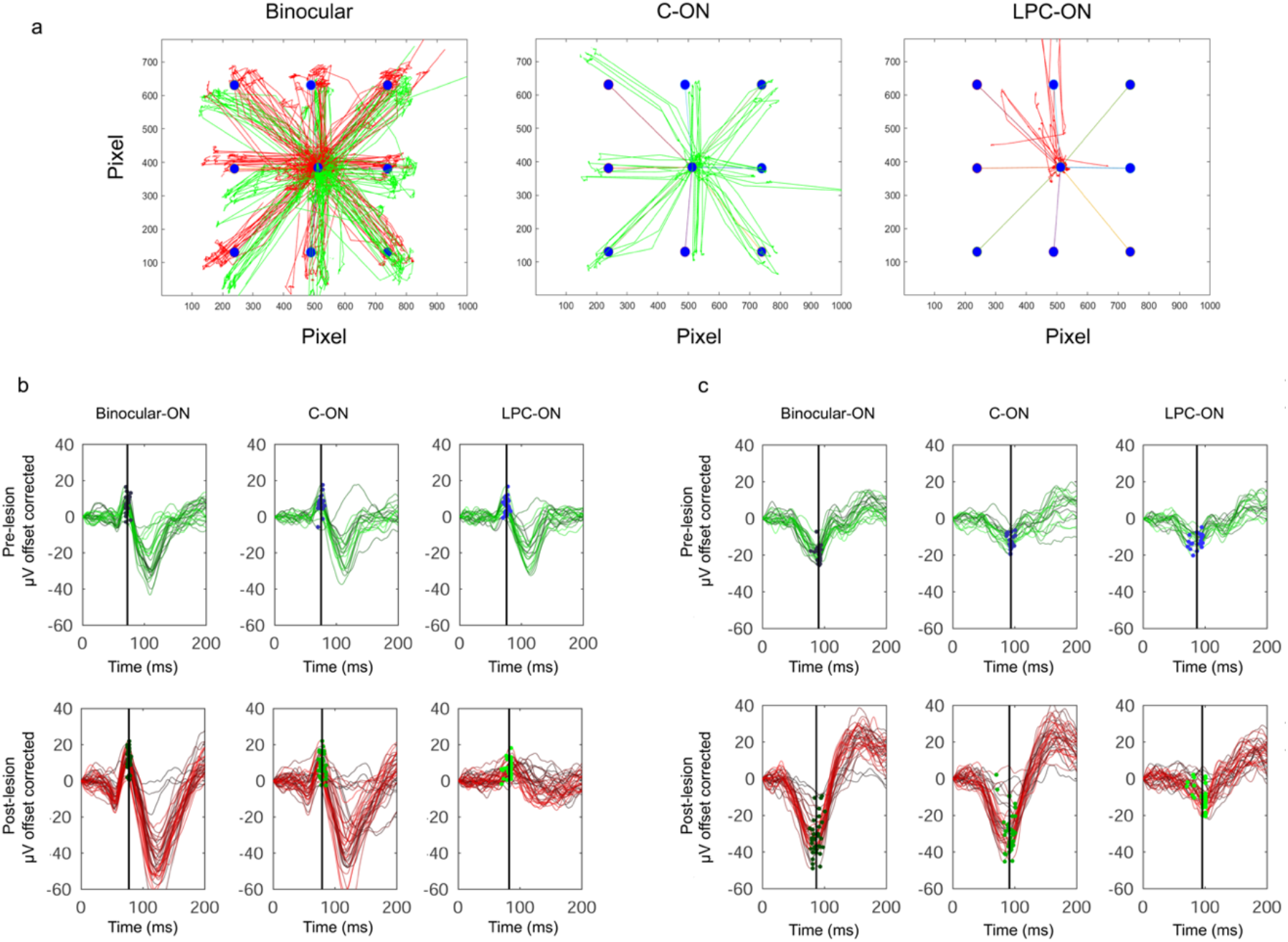
**(A)** Representative example of the cumulated latencies of eye saccades toward targets (black dots) presented randomly at central and eccentric locations of the screen in binocular and monocular condition. **(B and C)** Representative examples of **(B)** D30 checkerboard latency curves and **(C)** blue flashes, illustrating P80 **(B)** and N2 **(C)** peaks in the binocular and monocular conditions for one subject followed over 9 months.

**Fig. S2.**
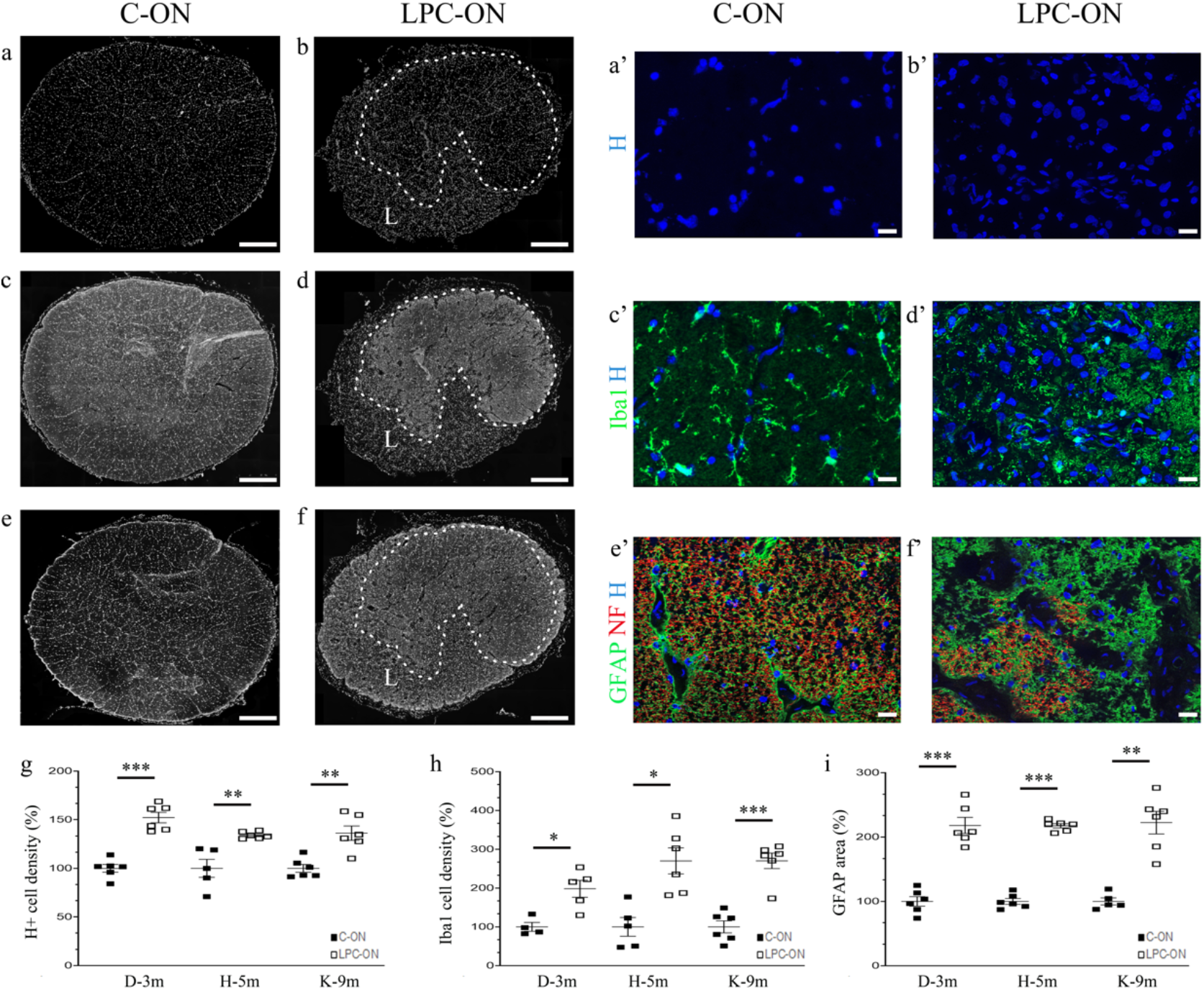
Immunohistochemical analysis of global cellularity, microglial and astroglial densities in CON **(A, A’, C, C, E, E’)** and LPC-ON **(B, B’, D, D’, F, F’). (A, A’, and B, B’**) The increased global cellularity evidenced by Hoechst staining in the lesion (L) is due mainly to the increased Iba1 **(C, C’, and D, D’)** and GFAP **(E, E’ and F, F’)** immunoreactivities. **(A’-F’)** are enlargements of **(A-F)**. Dotted lines in **B, D, F** represent the limits between the lesion and the spared parenchyma. (**G-I**) Quantification indicating the increase in global cellularity in **(G)**, Iba1+ **(H)**, and GFAP+ areas **(I)** at 3-, 5- and 9-months post-LPC. Data for each subject are expressed as a percentage of the C-ON fellow nerve values (n=1 per time-point). Graphs represent mean ± SEM, t-Test ** *P*<0.01, *** *P*<0.001. Scale bar **(A-F)** 200μm, **(A’-F’)** 15μm.

**Fig. S3.**
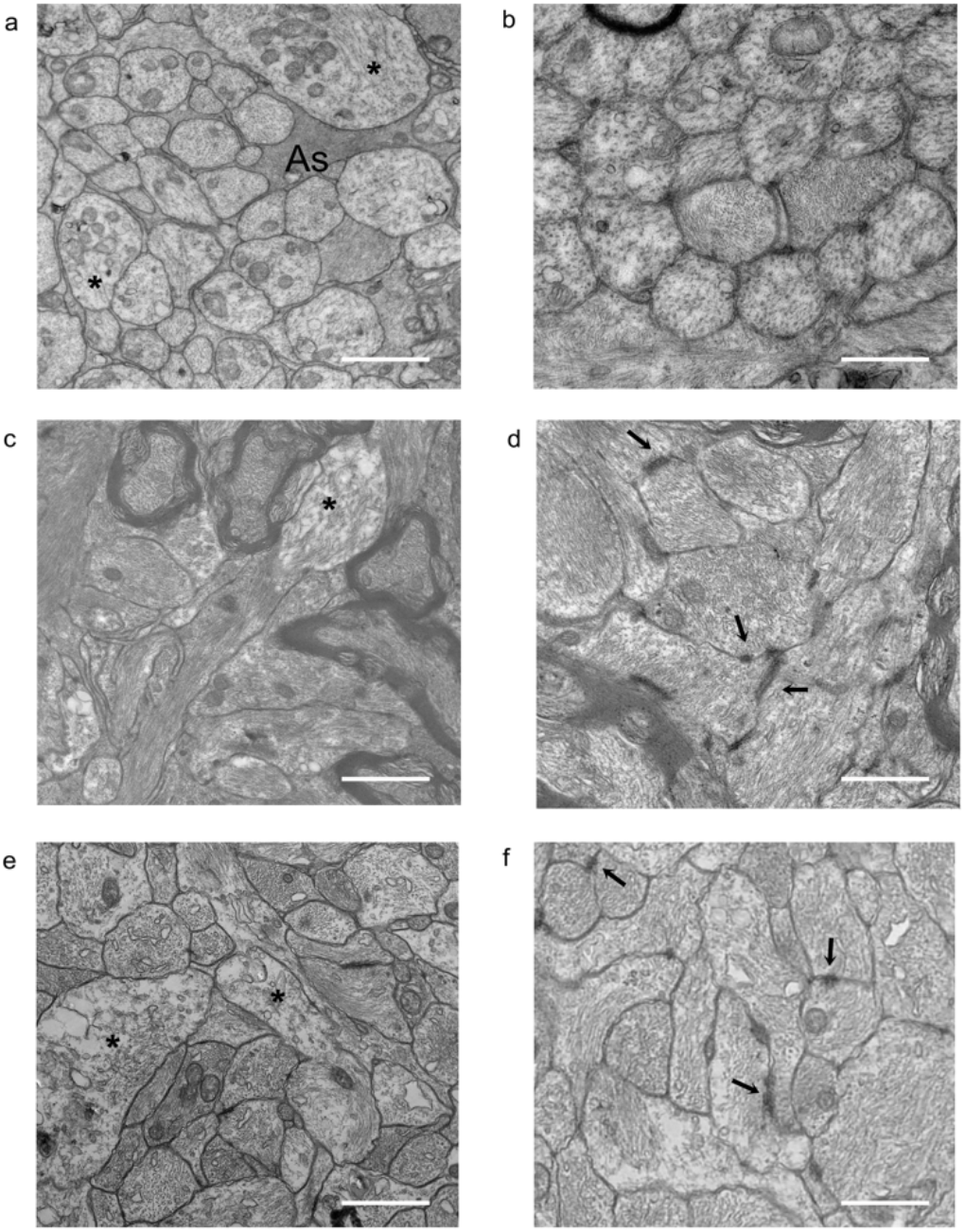
Ultrastructural evidence of axon modifications and increased axon-glial densities within the lesion. **(A, C, E)** center of the lesion, **(B, D, F)** periphery of the lesion. **(A, B)** At 3 months post-LPC, aggregated axons are surrounded by astrocyte processes (As). Axons contain neurofilaments and microtubules but some have increased mitochondrial contents (*)**. (C, D)** at 5- and **(E, F)** 9 months post-LPC, axons show swellings (*), increased neurofilaments but fewer microtubules. Arrows point to several areas of increased membrane densities. Scale bar **(A, B)** 2μm, **(C, D)** 1.6μm, **(E, F)** 1.2μm.

**Fig. S4.**
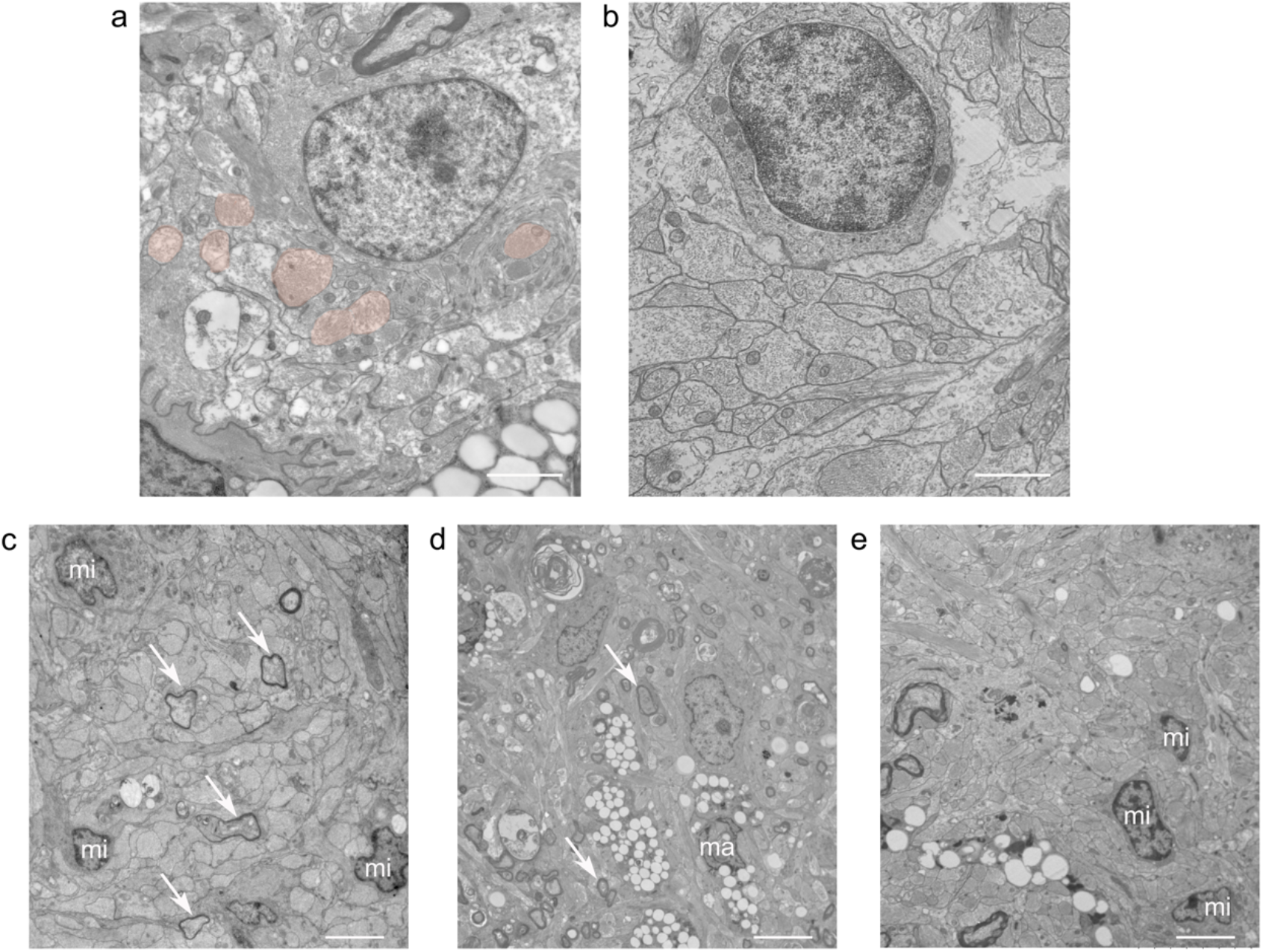
Electron microscopy evidence for long-term oligodendrocyte survival **(A, B)** and persisting inflammatory cells **(C-E). (A)** Oligodendrocyte in the lesion rim is in contact with-but does not form myelin around surrounding axons. **(B)** Oligodendrocytes in the lesion center surrounded by swollen astroglial processes. **(C-E)** Microglia (mi) and lipid-laden macrophages (ma) at the lesion rim at **(C**) 3, **(D)** 5-, and **(E)** 9 months post-LPC. White arrows indicate the presence of newly generated thin myelin sheaths. Scale bar **(A)** 2.5μm, **(B)** 1.4μm, **(C)** 3.5μm, **(D)** 7μm, **(E)** 4μm.

**Fig. S5.**
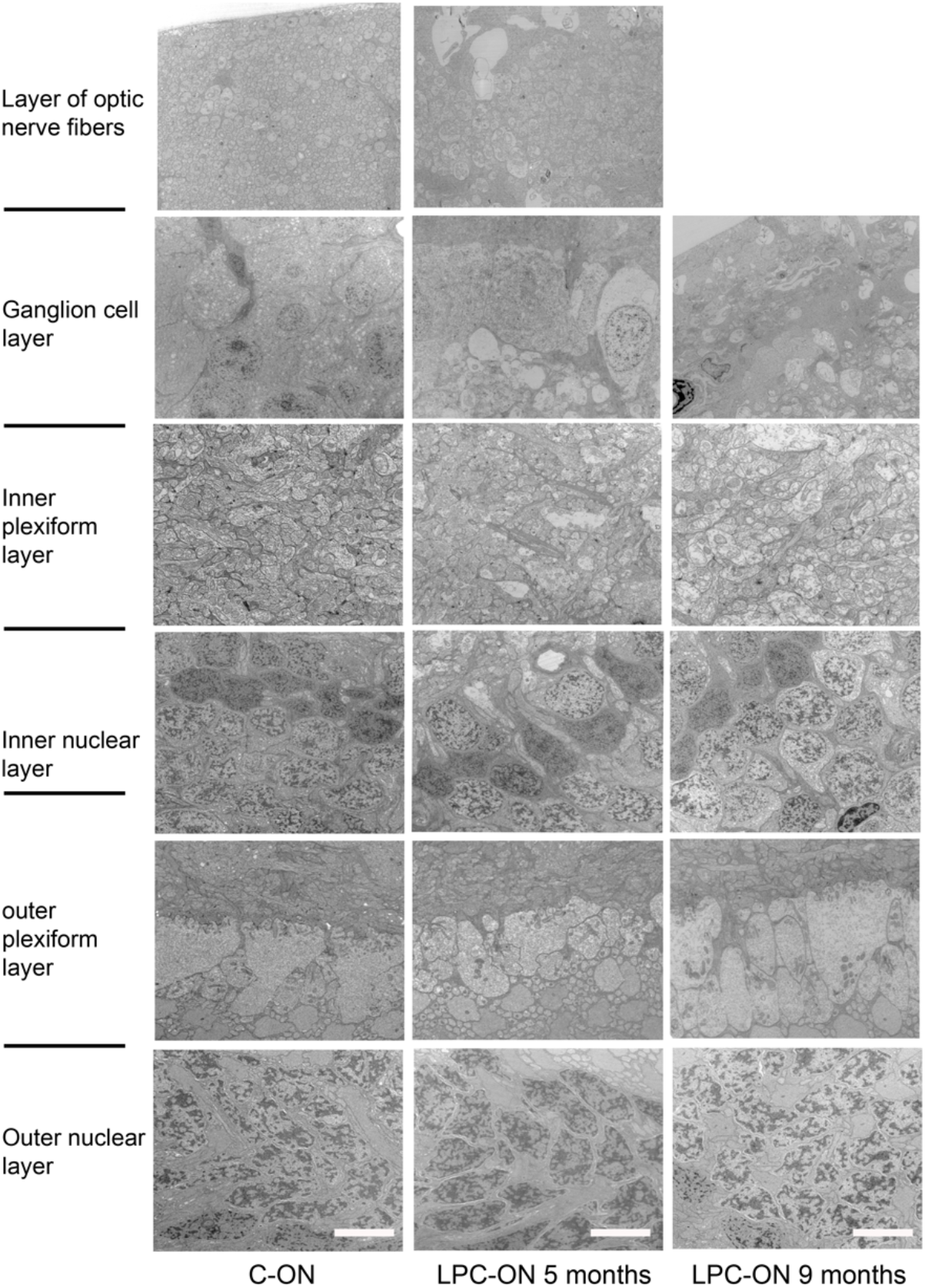
Ultrastructural characterization of the LPC-induced retinal modifications corresponding to C-ON (left) and LPC-ON (middle and right). Progressive degeneration of the optic nerve fiber layer at 5 months post-LPC, and complete disappearance by 9 months post-LPC. This is followed by modifications in the ganglion cell layer with vacuolation and disorganization, which are more severe at 9 months than 5 months post-LPC. Small axons of the inner plexiform layer contain multiple vacuoles. The outer plexiform and nuclear layers do not seem to be overly affected in the LPC-ON compared to the C-ON. Scale bar: 5μm.

**Fig. S6.**
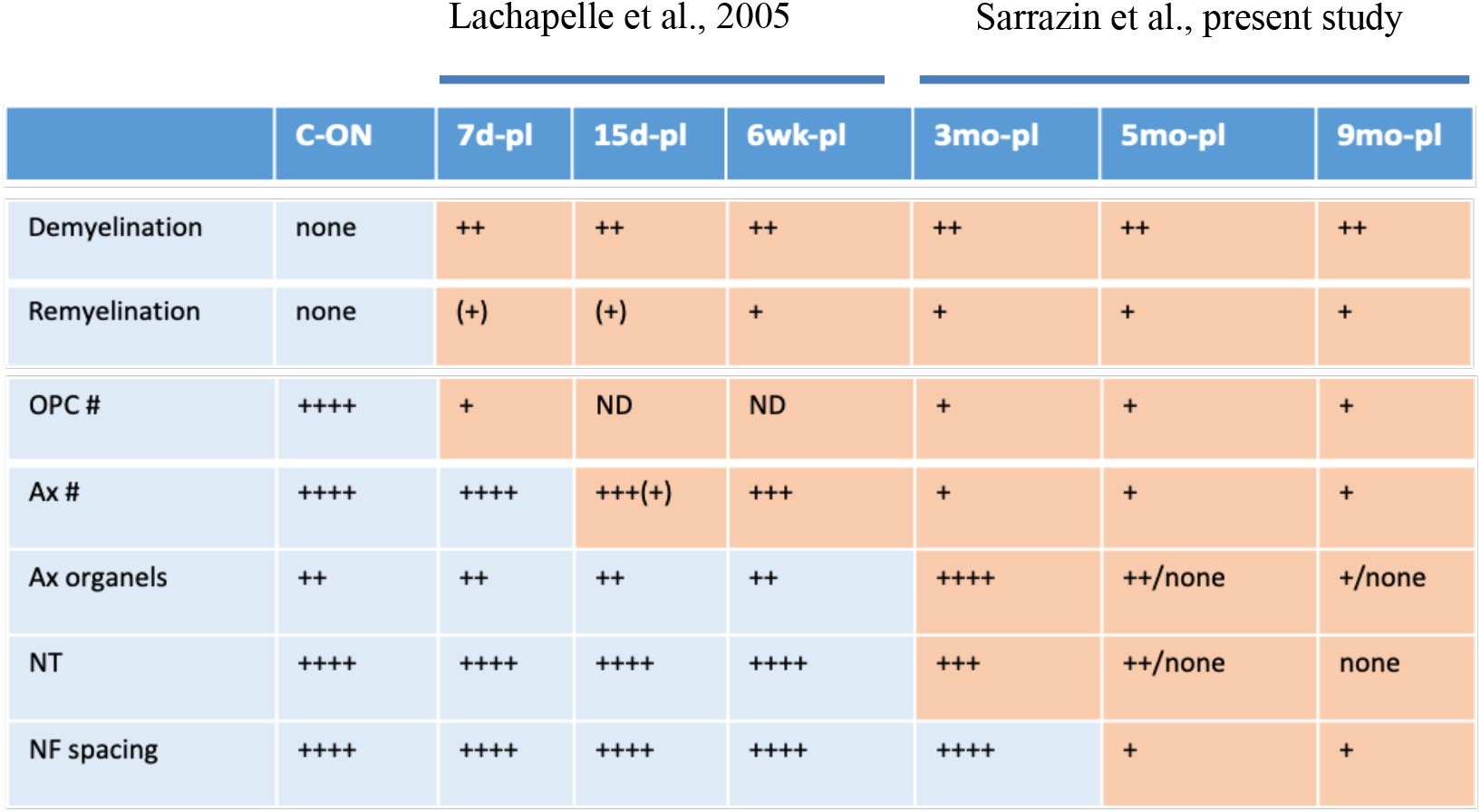
Summary of optic nerve temporal changes occurring in oligodendrocytes and axons following LPC-induced demyelination compared to the control fellow eye, and based on qualitative and quantitative data outlined in Lachapelle et al., 2005 (28) and the present study. Demyelination onsets early and remains steady followed by poor remyelination and decreased numbers of oligodendrocyte progenitors (OPC) and oligodendrocytes (OL). Axon (Ax) loss onsets slightly after 6 wk post-LPC and worsens from 3-9 months followed by altered axoplasmic organelles and decreased neurotubules (NT), and neurofilament (NF) spacing. The grey- and orange-colored areas highlight the normal-to-nearly normal- and pathological status of the nerve respectively. d:day, wk:week, mo:month, #:numbers.

**Table S1.** Experimental details.

**Table SI. Experimental details**
| Subjects | Post mortem analysis | OCT | ERG | VEP | PLR | VF | Time of sacrifice post-LPC |
| --- | --- | --- | --- | --- | --- | --- | --- |
| K-9m | IHC<br>-ON<br>-Ret | ND | Post-LPC | weekly | weekly | weekly | 9 months |
| Y-9m | ME<br>-ON<br>-Ret | ND | Post-LPC | weekly | weekly | weekly | 9 months |
| C-5m | EM<br>-ON<br>-Ret | ND | Pre -LPC<br>Post-LPC | weekly | weekly | weekly | 5 months |
| H-5m | IHC<br>-ON<br>-Ret | Post-LPC | Pre-LPC | weekly | weekly | weekly | 5 months |
| D-3m | IHC<br>-ON<br>-Ret | Post-LPC | Post-LPC | weekly | weekly | weekly | 3 months |
| G-3m | EM<br>-ON | ND | ND | ND | ND | ND | 3 months |

IHC = immunohistochemistry, EM = electron microscopy, Ret = retina, ND = not done,

**Table S2.**
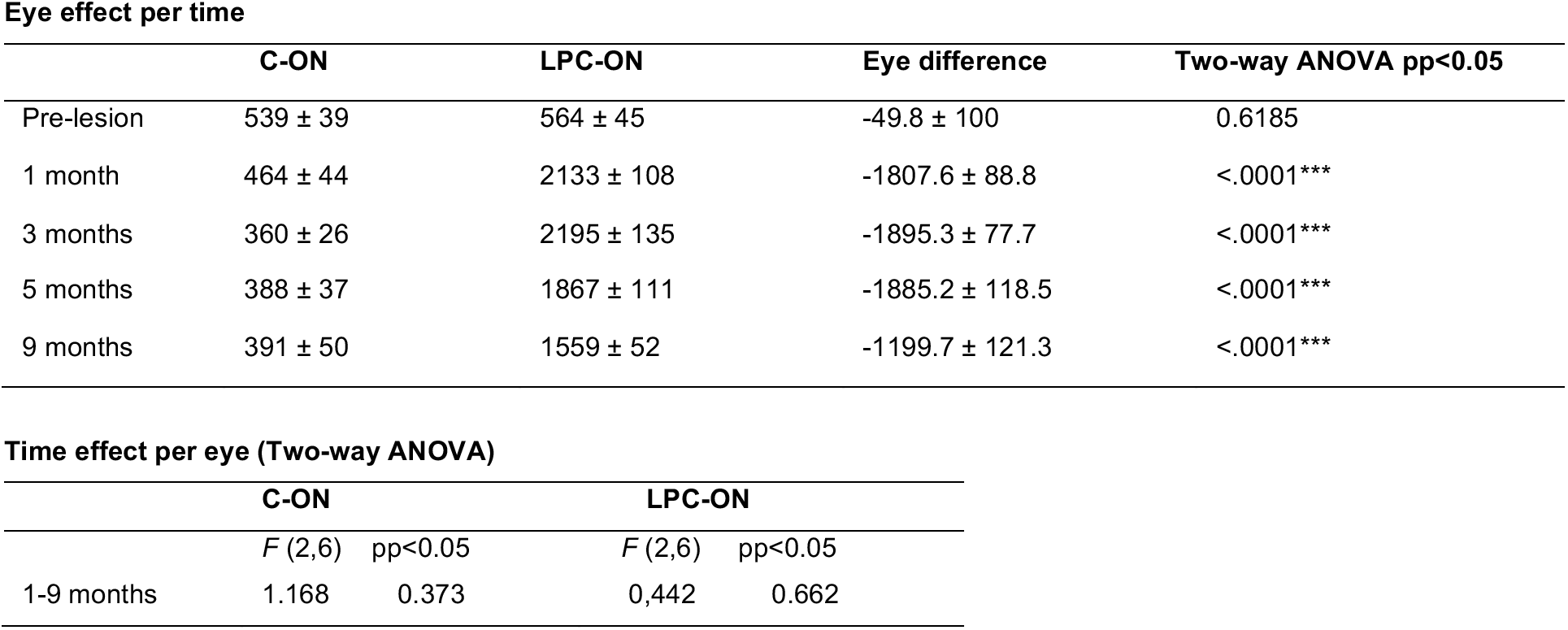
Follow-up of VF in 5 subjects (mean ± SEM)

**Table S3.** Follow-up of VEP latency in 5 subjects (mean ± SEM)

|  | C-ON | LPC-ON | Eye difference | Two-way ANOVA pp<0.05 |
| --- | --- | --- | --- | --- |
| Pre-lesion | 83.1 ± 0.9 | 83.1 ± 0.6 | - 0.065 ± 1.3 | 0.9603 |
| 1 month | 84.8 ± 0.7 | 96.0 ± 3.9 | - 11.165 ± 2.3 | <0.0001*** |
| 3 months | 83.0 ± 0.8 | 95.1 ± 2.8 | - 12.097 ± 2.5 | <0.0001*** |
| 5 months | 83.8 ± 1.7 | 97.1 ± 3.6 | - 13.329 ± 3.3 | <0.0001*** |
| 9 months | 85.9 ± 0.9 | 95.4 ± 2 | - 9.167 ± 3.2 | <0.0053** |

**Time effect per eye (Two-way ANOVA)**
|  | C-ON |  | LPC-ON |  |
| --- | --- | --- | --- | --- |
|  | *F* (3, 6) | pp <.05 | *F* (3, 6) | pp <.05 |
| 1-9 months | 0.489 | 0.702 | 4.513 | 0.056 |

**Checker board D30****Eye effect per time**
|  | C-ON | LPC-ON | Eye difference | Two-way ANOVA pp<0.05 |
| --- | --- | --- | --- | --- |
| Pre-lesion | 78.4 ± 0.7 | 78.5 ± 0.7 | - 0.123 ± 1.37 | 0.928 |
| 1 month | 80.3 ± 1.1 | 86.2 ± 3.3 | - 5.749 ± 2.43 | 0.0195* |
| 3 months | 79.2 ± 1.0 | 86.1 ± 4.3 | - 6.929 ± 2.46 | 0.0056** |
| 5 months | 76.4 ± 0.4 | 82.3 ± 3.6 | - 5.766 ± 3.60 | 0.1123 |
| 9 months | 75.5 ± 1.3 | 81 ± 1.9 | - 5.144 ± 3.12 | 0.1018 |

**Time effect per eye (Two-way ANOVA)**
|  | C-ON |  | LPC-ON |  |
| --- | --- | --- | --- | --- |
|  | *F* (3, 6) | pp<0.05 | *F* (3, 6) | pp <0.05 |
| 1-9 months | 2.913 | 0.123 | 1.064 | 0.432 |

**Red Flash****Eye effect per time**
|  | C-ON | LPC-ON | Eye difference | Two-way ANOVA pp <0.05 |
| --- | --- | --- | --- | --- |
| Pre-lesion | 81.5 ± 1.4 | 83.9 ± 0.9 | - 1.04 ± 1.3 | 0.4279 |
| 1 month | 86.7 ± 2.6 | 94.9 ± 3.9 | - 9.14 ± 2.9 | 0.0025** |
| 3 months | 81.1 ± 1.9 | 96.9 ± 4.9 | - 15.79 ± 2.7 | <0.0001*** |
| 5 months | 78.1 ± 5.3 | 95.2 ± 7.1 | - 17.14 ± 4.6 | 0.0003*** |
| 9 months | 80.7 ± 1.7 | 98.5 ± 5.3 | - 13.97 ± 5.2 | <0.0092** |

**Time effect per eye (Two-way ANOVA)**
|  | C-ON |  | LPC-ON |  |
| --- | --- | --- | --- | --- |
|  | *F* (1.15, 3.44) | pp <.05 | *F* (3, 9) | pp <.05 |
| 1-9 months | 2.571 | 0.199 | 3.888 | 0.049* |

**Blue Flash****Eye effect per time**
|  | **C-ON** | **LPC-ON** | **Eye difference** | **Two-way ANOVA pp<0.05** |
| --- | --- | --- | --- | --- |
| Pre-lesion | 83.6 ± 1.2 | 83.7 ± 1.2 | - 0.792 ± 1.41 | 0.5761 |
| 1 month | 86.1 ± 3.2 | 94.4 ± 3.9 | - 8.476 ± 3.33 | 0.0121* |
| 3 months | 84.4 ± 1.8 | 99.7 ± 3.0 | - 15.717 ± 2.99 | <0.0001*** |
| 5 months | 91.8 ± 6.1 | 96.4 ± 7.4 | - 4.570 ± 4.83 | 0.3462 |
| 9 months | 92.9 ± 3.2 | 93.4 ± 5 | - 1.344 ± 5.23 | 0.7976 |

**Time effect per eye (Two-way ANOVA)**
|  | **C-ON** |  | **LPC-ON** |  |
| --- | --- | --- | --- | --- |
|  | *F* (3, 9) | pp <0.05 | *F* (3, 9) | pp <0.05 |
| 1-9 months | 2.452 | 0.13 | 4.211 | 0.041* |

**Table S4.** Follow-up of PLR constriction amplitude (negative peak, C2) and postillumination pupillary reflex (PIPR), in 5 subjects (mean ± SEM)

**Eye effect per time**
|  | C-ON | LPC-ON | Eye difference | Two-way ANOVA pp<0.05 |
| --- | --- | --- | --- | --- |
| Pre-lesion | 56.7 ± 0.4 | 59.8 ± 0.7 | -3.12 ± 4.9 | 0.5343 |
| 1 month | 55.9 ± 1.5 | 86.7 ± 5.9 | -30.83 ± 3.8 | 2.962<sup>e</sup>-08*** |
| 3 months | 58.2 ± 0.8 | 85.4 ± 3.7 | -27.18 ± 3.8 | 2.560<sup>e</sup>-07*** |
| 5 months | 56.3 ± 2.2 | 78.6 ± 4.1 | -22.23 ± 4.3 | 2.641<sup>e</sup>-05*** |
| 9 months | 61.3 ± 1.9 | 77.8 ± 5.8 | -16.47 ± 6.1 | 0.0121* |

**Time effect per eye (Two-way ANOVA)**
|  | C-ON |  | LPC-ON |  |
| --- | --- | --- | --- | --- |
|  | *F* (3, 6) | pp <.05 | *F* (3, 6) | pp <.05 |
| 1-9 months | 1.408 | 0.282 | 2.072 | 0.162 |

**C2 Red Light** **Eye effect per time**
|  | C-ON | LPC-ON | Eye difference | Two-way ANOVA pp<0.05 |
| --- | --- | --- | --- | --- |
| Pre-lesion | 68.2 ± 0.6 | 64.7 ± 7.1 | 3.43 ± 4.2 | 0.4171 |
| 1 month | 61.4 ± 2.1 | 90.7 ± 3.5 | -29.21 ± 3.2 | 3.217e-09*** |
| 3 months | 64.2 ± 1.1 | 85.9 ± 3.6 | -21.69 ± 3.2 | 5.774<sup>e</sup>-07*** |
| 5 months | 63.3 ± 2.1 | 82.7 ± 2.3 | -19.35 ± 3.6 | 1.624<sup>e</sup>-05*** |
| 9 months | 69.9 ± 4.9 | 80.3 ± 1.5 | -10.38 ± 5.1 | 0.0527 |

**Time effect per eye (Two-way ANOVA)**
|  | C-ON |  | LPC-ON |  |
| --- | --- | --- | --- | --- |
|  | *F* (4) | *Pr* (>*F*) | *F* (4) | *Pr* (>*F*) |
| 1-9 months | 2.419 | 0.097 | 5.365 | 0.007** |

**PIPR Blue light** **Eye effect per time**
|  | C-ON | LPC-ON | Eye difference | Two-way ANOVA pp<0.05 |
| --- | --- | --- | --- | --- |
| Pre-lesion | 13.2 ± 2.6 | 12.3 ± 4.8 | 0.93 ± 4.5 | 0.8378 |
| 1 month | 22.6 ± 3.3 | 2.4 ± 2.2 | 20.21 ± 3.5 | 5.956<sup>e</sup>-06*** |
| 3 months | 14.6 ± 4.8 | 1.1 ± 1.8 | 13.53 ± 3.5 | 0.0007369*** |
| 5 months | 15.9 ± 3.7 | 3.2 ± 1.4 | 12.72 ± 3.9 | 0.0034073** |
| 9 months | 11.8 ± 0 | -1 ± 0.3 | 12.81 ± 5.5 | 0.0295* |

**Time effect per eye (Two-way ANOVA)**
|  | C-ON |  | LPC-ON |  |
| --- | --- | --- | --- | --- |
|  | *F* (4) | *Pr* (>*F*) | *F* (4) | *Pr* (>*F*) |
| 1-9 months | 1.117 | 0.387 | 3.346 | 0.040* |

**Table S5.** ERG (mean values ± S.D) of affected eye (LPC-ON) and fellow eye (C-ON) in 5 subjects.

|  | C-ON | LPC-ON |
| --- | --- | --- |
| Photopic flash (cone responses) - intensity: i-max # |  |  |
| a-wave amplitude | 33.5 ± 7.1 | 37.8 ± 6.7 |
| a-wave implicit time | 12.8 ± 1.5 | 12.8 ± 2.1 |
| b-wave amplitude | 115.0 ± 35.1 | 128.5 ± 40.0 |
| b-wave implicit time | 29.3 ± 0.5 | 29.5 ± 0.6 |
| Photopic flicker 30 Hz responses - intensity: i-max # |  |  |
| Fl-wave (30 Hz) | 110.9 ± 39.7 | 112.7 ± 26.9 |
| Scotopic (pure rod responses) dim flash intensity: -0.8 log cds/m2 |  |  |
| b-wave amplitude | 51.3 ± 21.2 | 59.8 ± 25.4 |
| b-wave implicit time | 51.3 ± 4.5 | 53.0 ± 2.9 |
| Scotopic (mixed rod-cone responses) intensity: i-max# |  |  |
| a-wave amplitude | 105.5 ± 22.6 | 101.3 ± 18.2 |
| a-wave implicit time | 14.0 ± 2.2 | 14.5 ± 1.3 |
| b-wave amplitude | 253.8 ± 66.4 | 256.0 ± 49.6 |
| b-wave implicit time | 35.8 ± 3.1 | 36.0 ± 2.6 |

i-max# = 1.0 log cds/m2
Amplitude values are in microvolts
Implicit time values are in milliseconds

**Table S6.**
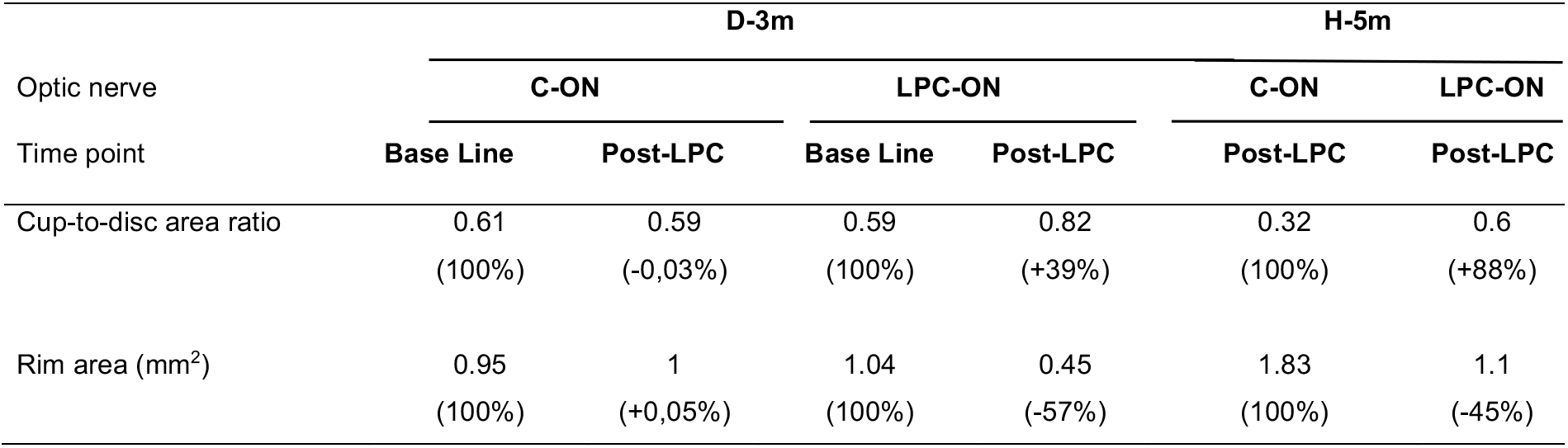
OCT assessment of the optic nerve head (ONH) parameters using the Glaucoma scan mode on two animals.

## References

1. Trapp BD, Nave K-A. Multiple Sclerosis: An Immune or Neurodegenerative Disorder? [Internet]. Annu. Rev. Neurosci. 2008;31(1):247–269.

2. Frohman E et al. Optical coherence tomography in multiple sclerosis. [Internet]. Lancet. Neurol. 2006;5(10):853–63.

3. Arnold DL. Changes observed in multiple sclerosis using magnetic resonance imaging reflect a focal pathology distributed along axonal pathways [Internet]. J. Neurol. 2005;252(S5):v25–v29.

4. Balk LJ et al. Timing of retinal neuronal and axonal loss in MS: a longitudinal OCT study. [Internet]. J. Neurol. 2016;263(7):1323–31.

5. Ortiz-Perez S et al. Visual field impairment captures disease burden in multiple sclerosis. [Internet]. J. Neurol. 2016;263(4):695–702.

6. de Seze J et al. Pupillary disturbances in multiple sclerosis: correlation with MRI findings. [Internet]. J. Neurol. Sci. 2001;188(1–2):37–41.

7. Forooghian F et al. Electroretinographic abnormalities in multiple sclerosis: possible role for retinal autoantibodies. [Internet]. Doc. Ophthalmol. 2006;113(2):123–32.

8. Cordano C et al. pRNFL as a marker of disability worsening in the medium/long term in patients with MS [Internet]. Neurol. - Neuroimmunol. Neuroinflammation 2019;6(2):e533.

9. Brusa A, Jones SJ, Plant GT. Long-term remyelination after optic neuritis: A 2-year visual evoked potential and psychophysical serial study. [Internet]. Brain 2001;124(Pt 3):468–79.

10. Chatziralli IP, Moschos MM, Brouzas D, Kopsidas K, Ladas ID. Evaluation of retinal nerve fibre layer thickness and visual evoked potentials in optic neuritis associated with multiple sclerosis. [Internet]. Clin. Exp. Optom. 2012;95(2):223–8.

11. Green AJ et al. Clemastine fumarate as a remyelinating therapy for multiple sclerosis (ReBUILD): a randomised, controlled, double-blind, crossover trial. [Internet]. Lancet (London, England) 2017;390(10111):2481–2489.

12. Picaud S et al. The primate model for understanding and restoring vision [Internet]. Proc. Natl. Acad. Sci. 2019;116(52):26280–26287.

13. Chanoumidou K, Mozafari S, Baron-Van Evercooren A, Kuhlmann T. Stem cell derived oligodendrocytes to study myelin diseases. [Internet]. Glia 2020;68(4):705–720.

14. Mozafari S, Baron-Van Evercooren A. Human stem cell-derived oligodendrocytes: From humanized animal models to cell therapy in myelin diseases [Internet]. Semin. Cell Dev. Biol. [published online ahead of print: October 2020]; doi:10.1016/j.semcdb.2020.09.011

15. Franklin RJM, Ffrench-Constant C. Regenerating CNS myelin - from mechanisms to experimental medicines. [Internet]. Nat. Rev. Neurosci. 2017;18(12):753–769

16. Melchor GS, Khan T, Reger JF, Huang JK. Remyelination pharmacotherapy investigations highlight diverse mechanisms underlying multiple sclerosis progression. ACS Pharmacol Transl Sci. 2019 Nov 14;2(6):372–386.

17. Bove RM, Green AJ. Remyelination pharmacotherapies in multiple sclerosis. Neurotherapeutics. 2017 Oct;14(4):894–904

18. Kondo Y et al. Spontaneous optic nerve compression in the osteopetrotic (op/op) mouse: a novel model of myelination failure. [Internet]. J. Neurosci. 2013;33(8):3514–25.

19. Mozafari S, Sherafat MA, Javan M, Mirnajafi-Zadeh J, Tiraihi T. Visual evoked potentials and MBP gene expression imply endogenous myelin repair in adult rat optic nerve and chiasm following local lysolecithin induced demyelination [Internet]. Brain Res. 2010;1351:50–56.

20. Schirmer L et al. Oligodendrocyte-encoded Kir4.1 function is required for axonal integrity. [Internet]. Elife 2018;7. doi:10.7554/eLife.36428

21. You Y, Klistorner A, Thie J, Graham SL. Latency delay of visual evoked potential is a real measurement of demyelination in a rat model of optic neuritis. [Internet]. Invest. Ophthalmol. Vis. Sci. 2011;52(9):6911–8.

22. Heidari M et al. Evoked potentials as a biomarker of remyelination [Internet]. Proc. Natl. Acad. Sci. 2019;116(52):27074–27083.

23. Teixeira LBC et al. Modeling the Chronic Loss of Optic Nerve Axons and the Effects on the Retinal Nerve Fiber Layer Structure in Primary Disorder of Myelin [Internet]. Investig. Opthalmology Vis. Sci. 2016;57(11):4859.

24. Satarian L, Kiani S, Javan M, Baharvand H. Transplantation of human induced pluripotent stem cell-derived neural progenitors to rat crushed optic nerve. Cell J. 2012;

25. Dehghan S, Javan M, Pourabdolhossein F, Mirnajafi-Zadeh J, Baharvand H. Basic fibroblast growth factor potentiates myelin repair following induction of experimental demyelination in adult mouse optic chiasm and nerves. [Internet]. J. Mol. Neurosci. 2012;48(1):77–85.

26. Cruz-Herranz A et al. Monitoring retinal changes with optical coherence tomography predicts neuronal loss in experimental autoimmune encephalomyelitis. [Internet]. J. Neuroinflammation 2019;16(1):203.

27. Moshiri A et al. A nonhuman primate model of inherited retinal disease. [Internet]. J. Clin. Invest. 2019;129(2):863–874.

28. Lachapelle F et al. Failure of remyelination in the nonhuman primate optic nerve. [Internet]. Brain Pathol. 2005;15(3):198–207.

29. Duncan ID et al. The adult oligodendrocyte can participate in remyelination. [Internet]. Proc. Natl. Acad. Sci. U. S. A. 2018;115(50):E11807–E11816.

30. Jäkel S et al. Altered human oligodendrocyte heterogeneity in multiple sclerosis. [Internet]. Nature 2019;566(7745):543–547.

31. Bacmeister CM et al. Motor learning promotes remyelination via new and surviving oligodendrocytes. [Internet]. Nat. Neurosci. 2020;23(7):819–831.

32. Tomiyama Y et al. Measurement of Electroretinograms and Visually Evoked Potentials in Awake Moving Mice. [Internet]. PLoS One 2016;11(6):e0156927.

33. Sriram P et al. Relationship between optical coherence tomography and electrophysiology of the visual pathway in non-optic neuritis eyes of multiple sclerosis patients. [Internet]. PLoS One 2014;9(8):e102546.

34. Soffer D, Raine CS. Morphologic analysis of axo-glial membrane specializations in the demyelinated central nervous system. [Internet]. Brain Res. 1980;186(2):301–13.

35. Henrikson CK, Vaughn JE. Fine structural relationships between neurites and radial glial processes in developing mouse spinal cord. [Internet]. J. Neurocytol. 1974;3(6):659–75.

36. Raine CS. Membrane specialisations between demyelinated axons and astroglia in chronic EAE lesions and multiple sclerosis plaques. [Internet]. Nature 1978;275(5678):326–7.

37. Green AJ, McQuaid S, Hauser SL, Allen I V., Lyness R. Ocular pathology in multiple sclerosis: retinal atrophy and inflammation irrespective of disease duration. [Internet]. Brain 2010;133(Pt 6):1591–601.

38. Gabilondo I et al. Dynamics of retinal injury after acute optic neuritis. [Internet]. Ann. Neurol. 2015;77(3):517–28.

39. Syc SB et al. Optical coherence tomography segmentation reveals ganglion cell layer pathology after optic neuritis [Internet]. Brain 2012;135(2):521–533.

40. Papakostopoulos D, Fotiou F, Hart JC, Banerji NK. The electroretinogram in multiple sclerosis and demyelinating optic neuritis. [Internet]. Electroencephalogr. Clin. Neurophysiol. 1989;74(1):1–10.

41. Franklin RJ, Ffrench-Constant C. Remyelination in the CNS: from Biology to therapy. Nat Rev Neurosci. 2008 Nov;9(11):839–55.

42. Wolswijk G. Chronic stage multiple sclerosis lesions contain a relatively quiescent population of oligodendrocyte precursor cells. J Neurosci. 1998 Jan 15;18(2):601–9.

43. Chang A, Tourtellotte WW, Rudick R, Trapp BD. Premyelinating oligodendrocytes in chronic lesions of multiple sclerosis N Engl J Med. 2002 Jan 17;346(3):165–73.

44. Kuhlmann et al. Differentiation block of oligodendroglial progenitor cells as a cause for remyelination failure in chronic multiple sclerosis Brain 2008 Jul;131(Pt 7):1749–58

45 Keirstead HS, Levine JM, Blakemore WF. Response of the oligodendrocyte progenitor cell population (defined by NG2 labelling) to demyelination of the adult spinal cord. Glia. 1998 Feb 22(2):161–70.

46. Johnson ES, Ludwin SK. The demonstration of recurent demyelination and remyelination of axons in the central nervous system. Acta Neuropathol. 1981;53(2):93–8.

47. Ehrlich M et al. Rapid and efficient generation of oligodendrocytes from human induced pluripotent stem cells using transcription factors [Internet]. Proc. Natl. Acad. Sci. 2017;114(11):E2243–E2252.

48. Buchet D, Garcia C, Deboux C, Nait-Oumesmar B, Baron-Van Evercooren A. Human neural progenitors from different foetal forebrain regions remyelinate the adult mouse spinal cord. [Internet]. Brain 2011;134(Pt 4):1168–83.

49. Mozafari S et al. Multiple sclerosis iPS-derived oligodendroglia conserve their properties to functionally interact with axons and glia in vivo. Sci. Adv. [published online ahead of print: 2020]; 2021, 6, 49: abc6983

50. Morales Pantoja IE et al. iPSCs from people with MS can differentiate into oligodendrocytes in a homeostatic but not an inflammatory milieu [Internet]. PLoS One 2020;15(6):e0233980.

51. Starost L et al. Extrinsic immune cell-derived, but not intrinsic oligodendroglial factors contribute to oligodendroglial differentiation block in multiple sclerosis. [Internet]. Acta Neuropathol. 2020;140(5):715–736.

52. Kirby L et al. Oligodendrocyte precursor cells present antigen and are cytotoxic targets in inflammatory demyelination. [Internet]. Nat. Commun. 2019;10(1):3887.

53. El Behi M et al. Adaptive human immunity drives remyelination in a mouse model of demyelination. [Internet]. Brain 2017;140(4):967–980.

54. Heß K et al. Lesion stage-dependent causes for impaired remyelination in MS. [Internet]. Acta Neuropathol. 2020;140(3):359–375.

55. Lloyd AF et al. Central nervous system regeneration is driven by microglia necroptosis and repopulation. [Internet]. Nat. Neurosci. 2019;22(7):1046–1052.

56. Miron VE. Beyond immunomodulation: The regenerative role for regulatory T cells in central nervous system remyelination [Internet]. J. Cell Commun. Signal. 2017;11(2):191–192.

57. Dutta R, Trapp BD. Mechanisms of neuronal dysfunction and degenration in multiple sclerosis Prog Neurobiol. 2011 Jan;93(1):1–12.

58. Luchetti S et al. Progressive multiple sclerosis patients show substantial lesion activity that correlates with clinical disease severity and sex: a retrospective autopsy cohort analysis [Internet]. Acta Neuropathol. 2018;135(4):511–528.

59. Kuhlmann T, Lingfeld G, Bitsch A, Schuchardt J. Acute axonal damage in multiple sclerosis is most extensive in early disease …. Brain 2002;

60. Rosolen SG, Kolomiets B, Varela O, Picaud S. Retinal electrophysiology for toxicology studies: applications and limits of ERG in animals and ex vivo recordings. [Internet]. Exp. Toxicol. Pathol. 2008;60(1):17–32.

61. Park JC et al. Toward a clinical protocol for assessing rod, cone, and melanopsin contributions to the human pupil response. [Internet]. Invest. Ophthalmol. Vis. Sci. 2011;52(9):6624–35.

62. Kankipati L, Girkin CA, Gamlin PD. The post-illumination pupil response is reduced in glaucoma patients. [Internet]. Invest. Ophthalmol. Vis. Sci. 2011;52(5):2287–92.

63. Kothari R, Bokariya P, Singh S, Singh R. A Comprehensive Review on Methodologies Employed for Visual Evoked Potentials [Internet]. Scientifica (Cairo). 2016;2016:1–9.

64. Klistorner A, Crewther DP, Crewther SG. Temporal analysis of the chromatic flash VEP-separate colour and luminance contrast components. [Internet]. Vision Res. 1998;38(24):3979–4000.

65. Lorenceau J, Humbert R. A multipurpose software package for editing two-dimensional animated images [Internet]. Behav. Res. Methods, Instruments, Comput. 1990;22(5):453–465.

## References

1. Nadal-Nicolás FM et al. Brn3a as a marker of retinal ganglion cells: qualitative and quantitative time course studies in naive and optic nerve-injured retinas. [Internet]. Invest. Ophthalmol. Vis. Sci. 2009;50(8):3860–8.

2. Schindelin J et al. Fiji: an open-source platform for biological-image analysis. [Internet]. Nat. Methods 2012;9(7):676–82.

3. Ehrlich M et al. Rapid and efficient generation of oligodendrocytes from human induced pluripotent stem cells using transcription factors [Internet]. Proc. Natl. Acad. Sci. 2017;114(11):E2243–E2252.

4. Freund P, et al. Nogo-A-specific antibody treatment enhances sprouting and functional recovery after cervical lesion in adult primates. Nat Med. 2006 Jul;12(7):790–2.

5. Beaud ML, et al. Anti-Nogo-A antibody treatment does not prevent cell body shrinkage in the motor cortex in adult monkeys subjected to un-lateral cervical cord lesion. BMC Neurosci. 2008 Jan 14;9:5.

